# Customizable silicification of DNA origami nanostructures

**DOI:** 10.1101/2025.10.28.685138

**Authors:** Anna Baptist, Lasse Guericke, Philipp Mauker, Oliver Thorn-Seshold, Amelie Heuer-Jungemann

## Abstract

The silicification of DNA origami nanostructures offers a powerful strategy for enhancing their mechanical stability and resistivity against detrimental environmental conditions. In the past years, several studies have investigated different aspects of the silica coating procedure, leading to several different silicification protocols. Until now, the silica coating generally served as a protective layer or as the base for the further deposition of inorganic materials. However, it did not carry any additional functionality itself. Here, we present two different approaches for the customization of the silica coating on DNA origami nanostructures. Firstly, we developed a custom synthesized silica precursor carrying a fluorescein molecule to endow the silica coating of both DNA origami monomers and crystals with fluorescence and show the applicability of this novel silicification for the stabilization and tracking of DNA origami nanostructures intracellularly. Secondly, we employ a silica precursor containing a disulfide bridge to develop a silica coating that is dissolvable in a reducing environment. We anticipate that the results presented in this study will expand the toolbox of silicification in DNA nanotechnology and will further pave the way towards applications in drug delivery and material science.

## Introduction

Since its invention in 2006, the DNA origami technique^1^ has become a versatile platform for a wide range of applications. It offers a highly programmable shape and a unique addressability for nm-precise modifications with guest molecules such as fluorophores, metal nanoparticles or proteins. Among other potential fields of use, DNA origami nanostructures have been envisioned as templates for the growth of complex hybrid materials in material science or as carrier structures for drug delivery.^2, 3^ In the past years, a variety of different DNA origami designs have been developed and studied with the goal of presenting proteins in a specific arrangement to cells or carrying a cargo designed to be released upon a particular stimulus.^4–10^ Furthermore, many different studies have already explored the distribution of DNA origami in cells (*in vitro*)^11^ or even in living organisms (*in vivo*)^12^. However, there are several challenges associated with the future application of DNA origami as a drug delivery system. Apart from considerations such as efficient targeting or controlled release mechanisms, a major obstacle is the lack of stability of DNA origami in biologically relevant conditions where DNases and low salt condition lead to the destabilization and degradation of the structures.

Diverse approaches to increase the stability of DNA origami against detrimental environmental influences have been presented, including UV point-welding, enzymatic ligation, or different types of protective coatings such as virus capsid proteins, oligolysine-PEG, or silica.^13–19^ Recently, the silicification of DNA origami (in solution or on surfaces, with coating thicknesses from sub-nm to several nm) has become an established and efficient method to increase structures’ chemical and biological stability against a variety of environmental cues^20–25^, making them more suited for materials and theranostic applications. Silicification also preserves their designed shapes and precise addressability, since single-stranded DNA (ssDNA) retains its hybridization ability even after silicification.^26^ While the shape and mechanical properties of the nanostructures will affect e.g. their cellular uptake behaviour^27–29^ as well as their ability to transport cargo, retaining addressability is of great importance for e.g. stimuli-responsive structures. Additionally, amorphous silica is generally considered to be biocompatible^30^, making silicified nanostructures and nanoparticles ideal candidates for biological applications.

However, while the addressability of DNA nanostructures has been maintained, the option to introduce further functionality into the silica coating has thus far not been explored. Here, we present two types of custom silica functionalisation for silicified DNA origami nanostructures, employing either fluorescent or redox-responsive silane precursors. When combined with the wide range of modifications possible on the DNA nanostructure itself, these offer a new and improved tool-kit for silicified DNA origami nanostructures for various purposes, including biomedical applications (**Figure 1)**.

**Figure 1:**
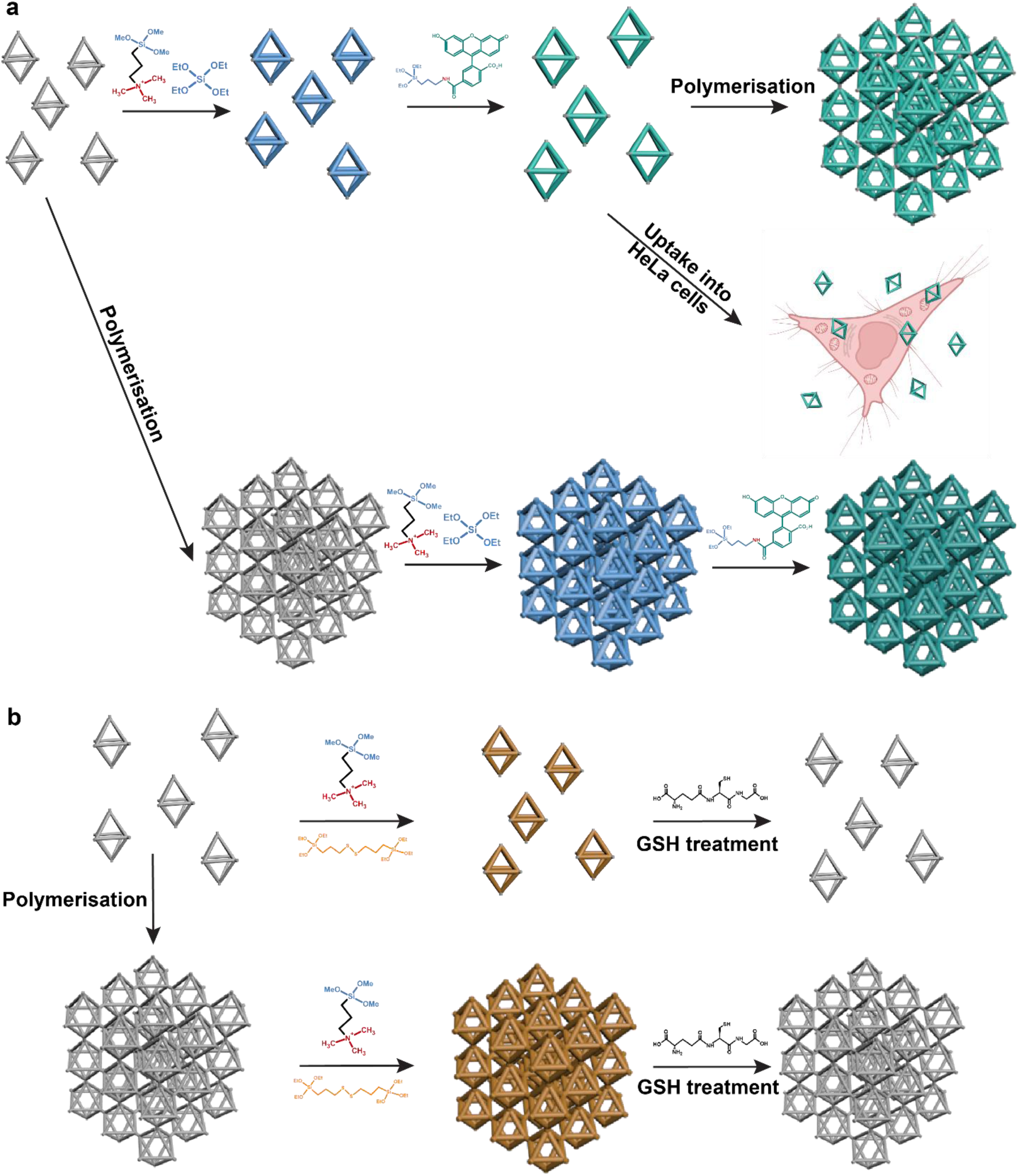
Schematic representation of the experimental workflow in this study. (a) Experiments with a fluorescent silane. (b) Experiments with a silica precursor containing a disulfide bridge.

***In our first customisation approach***, we introduce a fluorescein-labelled silane precursor, which can be incorporated during the silicification reaction, resulting in a fluorescent silica coating (**Figure 1a)**. Endowing nanostructures with fluorescence is of great interest for *in vitro* and *in vivo* applications: it has been very challenging to actually monitor the structural integrity of (bare or silicified) DNA origami nanostructures within cells, since the tracking of fluorescent modifications within the DNA origami only provides information about the presence of these fluorophores and their distribution in the sample - but not necessarily about the integrity of the entire nanostructure.^31, 32^ As shown by Lacroix et al., an intracellular fluorescent signal may well originate from dissociated fluorophores or partially degraded nanostructures.^33^ Other recently developed techniques such as origamiFISH^11^ are able to overcome these limitations to an extent. However, they are not easily applicable to DNA origami structures coated with an inorganic material such as silica. Therefore, endowing a silica coating with fluorescence would offer both enhanced stability of the coated DNA origami as well as the opportunity to monitor their structural integrity within a cellular environment. Different approaches for the creation of fluorescent silica nanoparticles have been presented, relying either on the covalent attachment of fluorescent dyes to silica precursors or on the physical entrapment of dyes into the pores or channels of a silica matrix.^34^ Nevertheless, the incorporation of fluorophores into the silica coating of silicified DNA nanostructures has not been shown to date. Here we show that by introducing a newly synthesized fluorescein-labelled silane precursor into the silicification reaction, a fluorescent silica coating can be integrated within single DNA origami monomers, as well as into large DNA origami crystals, where the fluorescence is distributed throughout the entire crystal. Despite the additional presence of many fluorophores in the silica, the stability of the structures in cell culture medium or in an intracellular environment is similar to that of standard silica coatings.

***In our second customisation approach***, we introduce the disulfide-containing silane precursor bis[3- (triethoxysilyl)propyl]disulfide (BTDS), which incorporates during silicification to give a DNA origami whose silica shell can be selectively disrupted by a reducing agent such as glutathione (GSH) (**Figure 1b**). This responsiveness of the silica coating to external stimuli can be highly attractive for drug release applications, or for controllably-degradable nanostructures that may alleviate concerns about potentially toxic accumulation of nanostructures in animals.^35–37^ For inorganic nanoparticles, different types of stimuli-responsive shells have long been conceived and demonstrated, e.g. nitrobenzyl-containing coatings inferring responsiveness to UV light.^38^ Among different responsive coatings, the integration of disulfide bridges is particularly attractive with regard to intracellular applications as they can easily be reduced in the presence of a reducing agent like GSH, which is present in cells in millimolar concentrations as a crucial antioxidant. Particularly cancer cells have been reported to exhibit elevated GSH levels.^39–41^ Very recently, an oligolysine-PEG coating for DNA origami was presented that contains disulfide bridges within the oligolysine segments, enabling it to undergo decomplexation in reductive environments.^42^ Furthermore, the use of disulfide bridge-containing silica precursors such as bis[3-(triethoxysilyl)propyl]tetrasulfide (BTSPTS) has been experimentally demonstrated for silica nanoparticles^43, 44^, showing that the nanoparticles could be dissolved upon contact with a reducing agent. Additionally, the coating of small DNA tetrahedra with a mixture of tetraethylorthosilicate (TEOS) and BTDS was carried out to achieve hybrid nanoframeworks that can efficiently deliver cargo such as doxorubicin intracellularly due to the dissolution of the silica shell in acidic and GSH-rich conditions.^45^ However, so far, such thiol-responsive functionalisation of the silica coating for DNA origami nanostructures has not yet been demonstrated; and stably incorporating the function-carrying silica reagents into coatings with thicknesses ranging from below one to several nm without negatively affecting the protective effect of the coating had remained an unsolved challenge.

## Results and discussion

### Fluorescent Silica Coatings

Generally, the silicification of DNA origami relies on two main silica precursors. Initially, positively charged trimethyl[3-(trimethoxysilyl)propyl]ammonium chloride (TMAPS) attaches electrostatically to the negatively charged DNA phosphate backbone, while the second precursor (usually tetraethylorthosilicate (TEOS)) then forms a covalent silica network around the DNA origami nanostructure^19^, not only forming an outer shell but also penetrating the internal structure of the DNA origami.^20^ In order to introduce fluorescence into the silica coating, we therefore synthesised a fluorescein-conjugated silane monomer, which could be added to the reaction mixture; for this we acylated carboxyfluorescein with (triethoxysilyl)propylamine, expecting the amide product to be chemically stable during silicification (see Experimental Methods and **Figure S1**). Due to their versatility, we chose the well-studied DNA origami octahedra for all silicification experiments (see **Figure S2** for the design). Importantly, these octahedral monomers can be polymerised into cubic crystals via sticky end hybridization, even after silicification of the monomers as we were able to show in a previous study^26^. Experiments were performed on DNA origami monomers with an ultrathin silica coating and subsequently assembled crystals as well as on pre-formed crystals that were silicified post-assembly. While monomers were used for biological tests (stability in cell culture medium and *in cellulo*), the DNA origami crystals allowed to study the distribution of the fluorescence inside such large structures.

Since the newly synthesized fluorescent silane lacks one binding site compared to TEOS, we hypothesised that it may cause an increased porosity of the silica network. In addition, we anticipated that the larger size of the fluorescein sidechain compared to the usual fourth ethoxy group might decelerate the reaction. Therefore, a ‘two-step-silicification’ approach was devised (**Figure 2a**). This procedure includes a first silicification step using the standard precursors TMAPS and TEOS to initially create a thin, stable silica coating around the DNA origami, followed by a purification step to remove excess precursor molecules, and subsequently a second much longer silicification step using the fluorescent silane to create a second, but fluorescent silica layer. This was again followed by another purification step to remove the unbound fluorescent silane.

**Figure 2:**
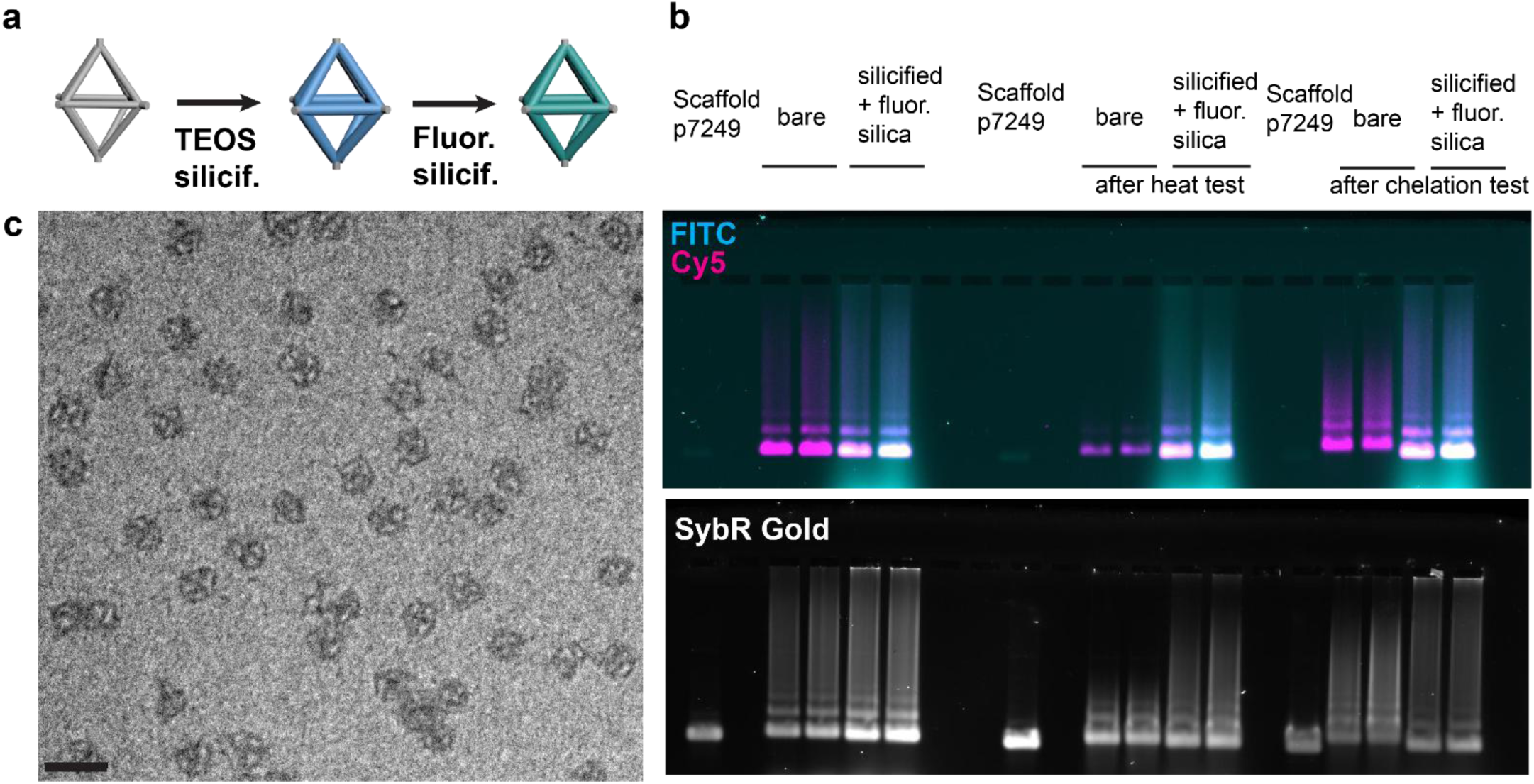
(a) Scheme depicting the experimental workflow for the fluorescent silicification of octahedral DNA origami monomers. (b) Agarose gel electrophoresis showing the successful integration of the fluorescent silane into the silica coating of the octahedral monomers and the enhanced stability of the silicified structures compared to bare octahedra. (c) TEM image of fluorescently silicified octahedra after a heat test. Scale bar: 100 nm.

In order to allow for identification of both the DNA origami and the silica coating by fluorescence, a set of Cy5-labelled staples was incorporated into the structures which were then analysed using agarose gel electrophoresis (AGE). As can be seen from **Figures 2b** and **S3**, the fluorescent precursor was successfully incorporated into the silica coating and no significant unspecific binding of the fluorescent silane or the free fluorescein dye to either bare or pre-silicified DNA origami could be observed. Having established successful incorporation of the fluorescent silane, we next tested the stability of the resulting structures with respect to heat and low salt conditions using AGE and TEM (see also **Figure 2c** for a TEM image of heat-tested octahedra). As expected, the bare octahedra clearly lost their structural integrity when subjected to the detrimental conditions, indicated by the upward shift in the gel bands and the loss in signal of the internal fluorescent modification (Cy5 channel). The fluorescently silicified nanostructures, however, did not show signs of degradation and continued to exhibit a strong signal from both the internal fluorophores and the fluorescein-containing silica coating, proving the increased stability inferred by the silicification (**Figure 2b**).

Since AGE displayed good co-localization of fluorescein and Cy5, we next made use of the ability of the silicified octahedra to form micron-sized cubic crystals to visualise the successful incorporation of the fluorescent silane by confocal microscopy (**Figure 3a**). For crystal formation, fluorescently silicified octahedra with two different types of sticky ends (type A and B) were mixed and polymerised via a heat ramp. The resulting cubic crystals were then attached to a microscopy slide and imaged via confocal microscopy, showing strong fluorescent signal in both FITC and Cy5 channels (**Figure 3b**). Importantly, this also successfully showed that the fluorescent silane remained incorporated even after undergoing a heat ramp for several days for crystal formation.

**Figure 3:**
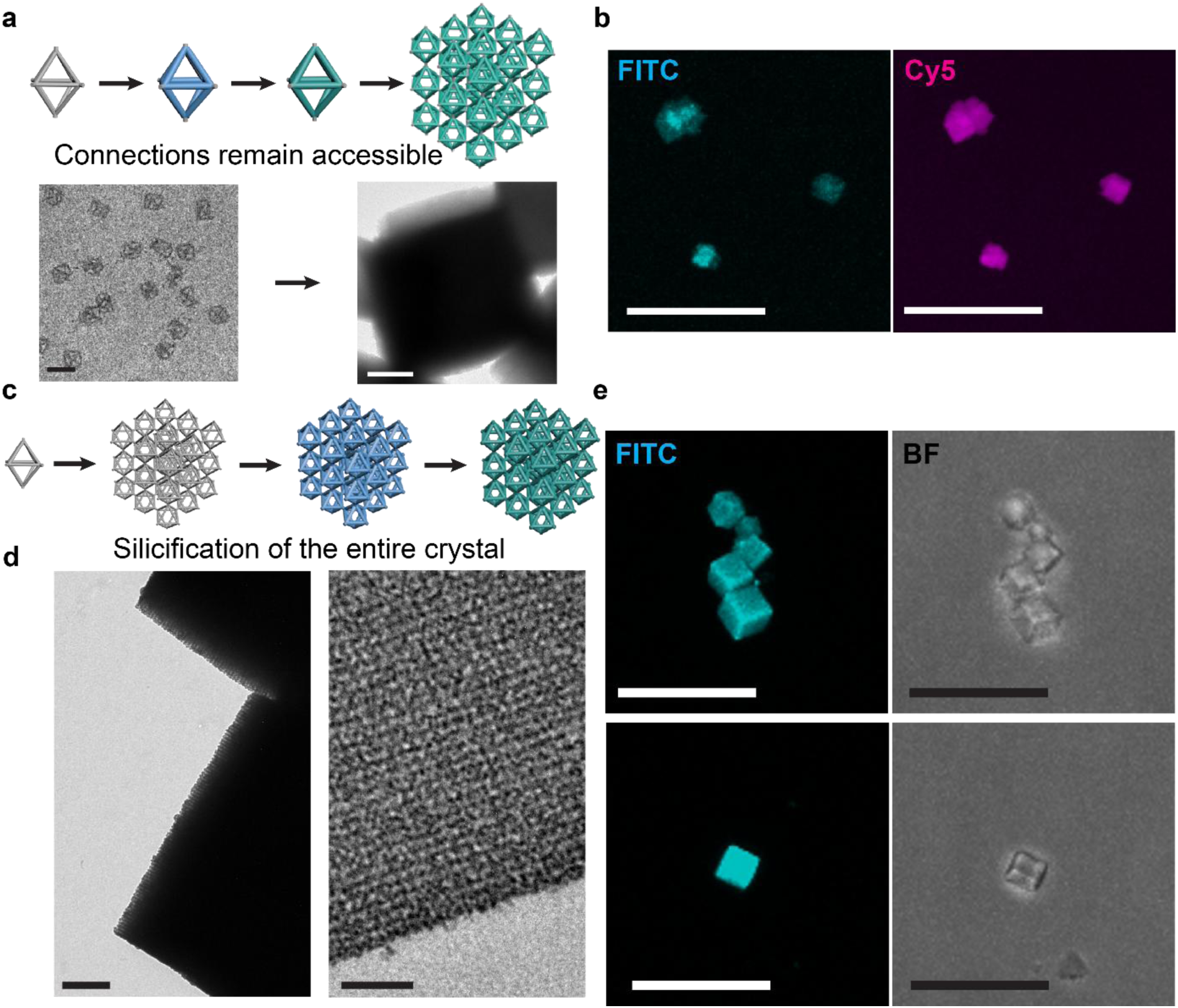
Fluorescent silicification of DNA origami crystal structures. (a) and (c) Schematic representation of the respective experimental workflow for the creation of silicified DNA origami crystals. (b) Confocal microscopy images of crystals obtained from the polymerisation of fluorescently silicified octahedra. Scale bars: 20 µm. (d) 3D projections obtained from z-stacks recorded using confocal microscopy. The imaged fluorescently silicified DNA origami crystals show a strong signal in the FITC channel, indicating the successful integration of a relatively large amount of fluorescent silica. Scale bars: 2 µm. (e) Left: TEM image showing a close-up of the fluorescently silicified crystals. The strong contrast in the images is indicative of a thick silica coating. Right: Crystal slice showing the internal structure of the fluorescently silicified DNA origami crystals. The order within the crystals is maintained and the crystals appear to be homogeneously silicified. Scale bars: 20 µm.

Thus far we only showed that the fluorescent silane could be successfully incorporated into ultrathin silica coatings. To investigate the incorporation into much thicker coatings, we next used pre-formed DNA origami crystals for silicification. For this, the cubic crystals created from the octahedral monomers were silicified post-assembly to create a thick silica coating (**Figure 3c**). In order to determine the optimal protocol for the creation of a strongly fluorescent silica layer, several different timepoints throughout the silicification procedure were tested for the addition of the fluorescent silane to the crystals. We thus found that the addition of the fluorescent silane after 8-10 h of silicification with a subsequent overnight incubation led to the best results, namely a well-distributed fluorescein signal (not only on the edges of the crystals) and a fully fluorescent cubic crystal structure. Z-scans through the entire crystals and the 3D projections thereof showed a strong fluorescein signal from all crystal slices in the microscopy images (**Figures 3d** and **S4, Supplementary Videos**). TEM micrographs confirmed that the silicification resulted in a comparatively thick and uniform silica coating, yielding highly stable crystals (upon exposure to cell culture medium, elevated temperatures and chelation conditions) with fully retained lattice order (**Figures 3e** and **Figure S5**).

It is important to note that the time point of the addition of the fluorescent silane had a strong effect on the experimental results. Adding the fluorescein-modified precursor shortly after the beginning of the silicification procedure led to an incomplete coverage of the crystals which only exhibited a fluorescent signal on the outer periphery of the cubes and whose surfaces appear to be irregular and bulky (**Figure S6**). We hypothesize that the early addition of the precursor lead to the increased formation of silica clusters, resulting in the inhomogeneous coating appearance.

After having established the proper incorporation of the fluorescent silane, we next sought to test their behaviour and stability of fluorescently silicified DNA origami under biological conditions (exposure to cell culture media and intracellular stability). It has been previously observed for silica nanoparticles that certain components of cell culture media can etch the silica layers within hours, causing dents in the silica surface and (partially) exposing core materials. This erosion of the nanoparticles was mostly attributed to the abundance of amino groups in the cell culture medium. However, it was also found that a higher level of polymerisation within the silica network enhances the stability.^46, 47^ In light of these observations, it is of high interest to investigate the stability of the silica coating itself in cellular environments. To study the stability of our structures and be able to compare bare and (fluorescently) silicified origami, all DNA origami structures without the fluorescent silane were equipped with both Cy5- and FITC-modified staples, whereas the fluorescently silicified samples were equipped with Cy5-carrying staples only. The incorporation of two different internal fluorescent modifications into the (bare) DNA origami samples and the monitoring of the co-localisation of these two signals can also serve as a tool for intracellular localisation and allow for a qualitative assessment of their structural integrity based on co-localisation of the two dyes.

Therefore, to investigate the stability of our structures in biologically relevant conditions, we incubated the structures (bare, silicified and fluorescently silicified) in FluoroBrite DMEM cell culture medium with a final concentration of 10% fetal bovine serum (FBS) for up to 24 h (**Figure S7**). To ensure the full stability not only of the DNA origami within the silica coating, but also of the fluorescent outer silica layer, the reaction time with the fluorescent silane was significantly prolonged compared to the duration of a ‘standard silicification’ (see Experimental Methods and **Figure S8**). While the bare DNA origami exhibited clear signs of disassembly already after 2 h, a significant fluorescein signal was retained in the fluorescently silicified sample even after 24 h, indicating that the prolonged incubation with the fluorescent silane during silicification led to a stable silica network and a sufficient incorporation of the fluorescent precursor. The silicified octahedra with internal fluorescent modifications were found to be overall stable and showed only a minor increase in unbound FITC-modified staples. Together with the results from cell culture medium tests of fluorescently silicified octahedra with only short incubation times with the fluorescent silane (**Figure S8**), where the structures appeared to maintain their shapes while a strong release of the fluorescein signal after 24 h was observed, our findings imply that our novel silica precursor requires significantly more time for the build-up of a reasonably stable silica layer than TEOS. This could on the one hand be attributed to the reduced number of reactive groups (three instead of four), which can participate in the formation of siloxane bridges, and on the other hand to the bulky fluorescein modification that could lead to steric hindrance and a reduced diffusion speed, resulting in a thinner, more porous fluorescent layer.

After having successfully established the stability of our fluorescently silicified structures in cell culture medium, we next sought to investigate their behaviour *in cellulo*. For this, bare and (fluorescently) silicified structures were incubated with HeLa cells and their uptake and intracellular stability were observed by confocal microscopy (see **Figure 4** and **Figure S9**).

**Figure 4:**
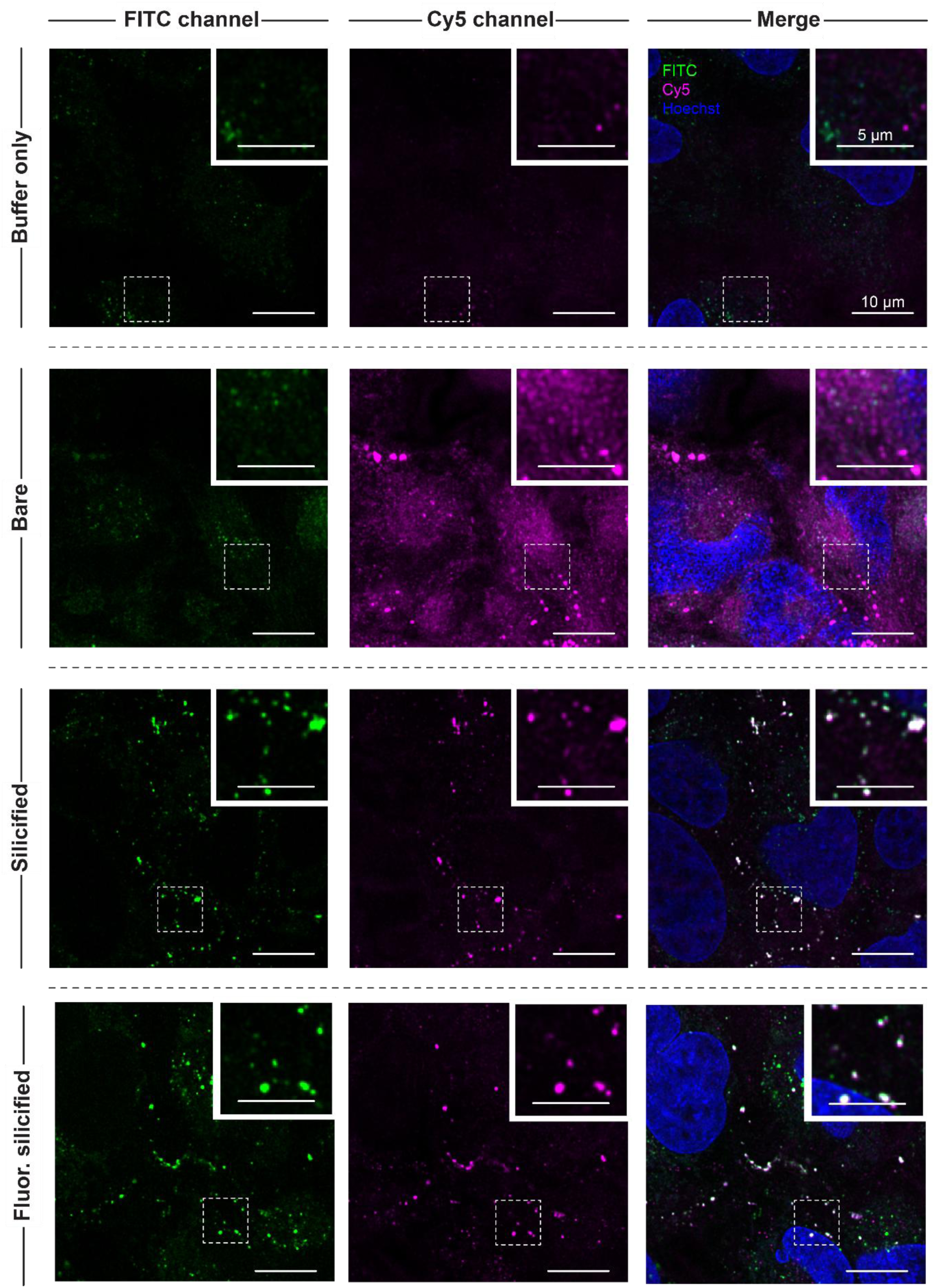
Cellular uptake of bare, silicified and fluorescently silicified DNA origami nanostructures. Representative images are shown after 6 h (bare octahedra) or 24 h (silicified octahedra) of incubation. Control samples (first row) incubated only with buffer for 24 h show only minor autofluorescence. Already after 6 h, the bare DNA origami nanostructures (second row) were clearly degraded. Both silicified and fluorescently silicified samples (third and fourth row) exhibited significant colocalisation between the fluorescein and the Cy5 signal, indicating the stability of the structures even after 24 h. Scale bars: 10 µm, insets: 5 µm.

As can be seen from the confocal images of the bare structures, already after 6 h, no significant co-localisation of the FITC and Cy5 signals could be observed; instead, particularly the Cy5 signal spread throughout the cells, strongly indicating a separation from the DNA origami. Contrary to that, the DNA origami with a standard silicification exhibited an excellent overlap of the two fluorescent signals even after 24 h. Even though the integrity of the octahedral shape cannot directly be monitored, the signal overlap indicates a strong stabilization of the monomers through the silicification. Encouragingly, also the fluorescently silicified samples exhibited a significant co-localisation of the Cy5 and the fluorescein signal inside the cells even after 24 h of incubation, suggesting that the fluorescent silica layer remains mostly intact. However, it has to be noted that some of the fluorescein signal did not overlap with any Cy5 signal.

This fluorescein signal was found mostly in a perinuclear localisation, suggesting lysosomal accumulation. To investigate if the signal could arise from insufficient purification from the fluorescein-silane precursor, we conducted control experiments using confocal microscopy where HeLa cells were incubated with the silane either directly from the stock solution or after a four-day incubation with TMAPS and TEOS. As can be seen from **Figure S10**, no significant intracellular fluorescent signal could be observed after incubation with the fluorescent silane only. However, the control sample from a ‘mock silicification’ (TMAPS + TEOS + fluorescent silane, without DNA origami) yielded several bright fluorescent spots in the FITC channel which can again mostly be found in a perinuclear localisation, very similar to the punctuated fluorescein-only signal observed in the samples with fluorescently silicified DNA origami. We therefore hypothesise that the fluorescein signal originates either from small silica aggregates formed during the silicification of the DNA origami, which could not be fully removed during the purification steps, or to a small extent from a partially etched outer silica layer from the DNA origami, in agreement with the medium tests (**Figure S7**). Nevertheless, for the fluorescently silicified DNA origami, the vast majority of Cy5 signal showed excellent co-localisation with the fluorescein signal, suggesting that structures remained intact and retained their fluorescent coating even after 24 h.

The experimental results here therefore establish our novel fluorescent silica precursor as an effective tool for the investigation of the intracellular distribution and stability of silicified DNA origami nanostructures.

### Degradable Silica Coating

After having successfully introduced fluorescence into the silica coating, we next investigated whether the silica coating could also be rendered degradable on demand. Particularly in the light of future *in vivo* applications of silicified-DNA origami nanostructures, the ability to create an on-demand degradable coating is highly desirable not only for drug delivery applications, but also to avoid potential accumulation in cells or certain organs resulting in toxicity. An interesting candidate for such a dissolvable silica layer is BTDS, a silica precursor, that resembles a dimer of TEOS molecules, but contains a disulfide bridge (see **Figure 1b**). Upon reduction of the disulfide bridge, a crack in the silica coating is formed, making the underlying DNA origami nanostructures again susceptible to environmental conditions. In order to create such a dissolvable silica coating, TEOS was replaced by BTDS during the silicification reaction. As preliminary tests revealed that the standard silicification time of several hours was insufficient to create a protective coating (see **Figure S11a** and **b**), the incubation time with both TMAPS and BTDS was extended to 4 d, followed by a purification step using spin filtration, resulting in a stable coating. We hypothesise that the presence of the disulfide bond and the larger size of the molecule resulted in a slower reaction rate, therefore requiring the longer reaction time, similar to the fluorescein silane precursor. In a next step, we tested the degradability of the silica coating. For this, a sample containing BTDS-silicified octahedra was incubated with a freshly prepared solution of GSH for 3 h. After a purification step to remove the GSH, the sample was subjected to both a chelation and a DNase I degradation test and subsequently analysed by AGE and TEM. As can be seen from **Figure 5**, BTDS-silicified structures that were not pre-treated with GSH successfully withstood the detrimental conditions and maintained their structural integrity both after chelation and DNase tests. However, if treated with GSH, structures were clearly degraded when subjected to chelating conditions or DNases, indicating successful reduction of the disulfide bridge by GSH and subsequent degradation of the silica coating. Interestingly, even though GSH did not affect the structural integrity of the bare DNA origami, the digestion of GSH-treated DNA origami was accelerated compared to untreated bare DNA origami (**Figure 5** and **Figure S12**).

**Figure 5:**
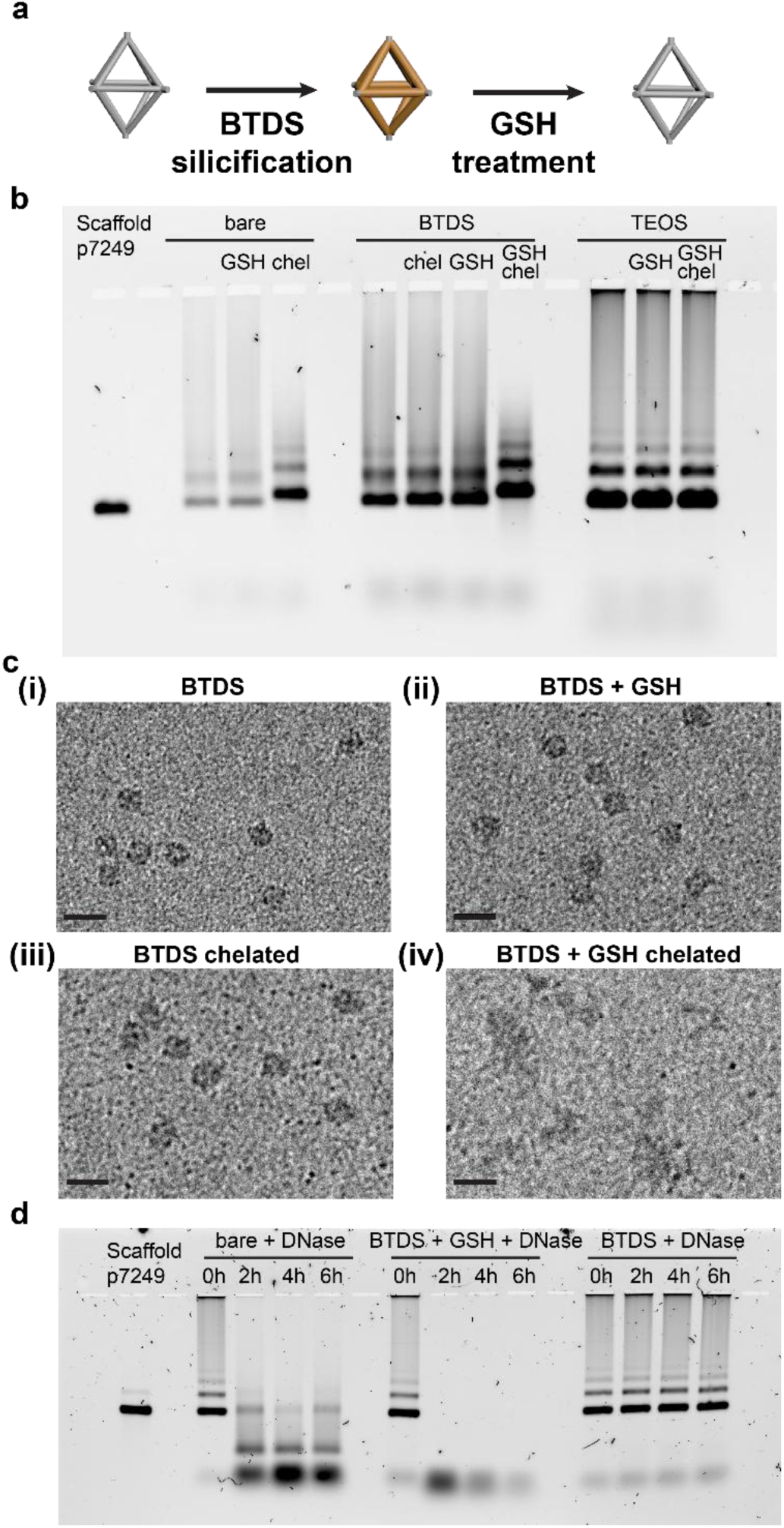
BTDS silicification of octahedral DNA origami monomers. (a) Scheme representing the BTDS silicification of octahedral monomers, followed by the incubation in GSH, leading to the dissolution of the silica coating. (b) Agarose gel showing the successful silicification of DNA origami octahedra with BTDS instead of TEOS. While the treatment with 10 mM GSH for 3 h does not seem to have a noticeable effect on samples with a standard silicification, the BTDS-silicified and GSH-treated sample clearly seems to be degraded during a chelation test, similar to a bare sample. (c) TEM images for (i) BTDS-silicified monomers directly after purification, (ii) BTDS-silicified monomers after treatment with GSH for 3 h, (iii) BTDS-silicified octahedra after incubation in 1x TAE buffer for 3 h, and (iv) BTDS-silicified and GSH-treated octahedra which are disassembled. Scale bars: 100 nm. (d) DNase test for different DNA origami samples. While the bare DNA origami octahedra (left) are rapidly degraded by the DNases, the BTDS-silicified monomers (right) can withstand even 24 h of incubation with the DNases. Interestingly, the BTDS-silicified samples after GSH treatment are exhibit an even faster and very complete digestion already within the first 2 h of incubation with the DNase.

All in all, our results show that the silicification with BTDS can indeed be used to create a silica coating that effectively shields the DNA origami nanostructures from detrimental conditions such as DNases and low salt environments, but that can also – unlike ‘standard’ silica coatings – be dissolved in the presence of a reducing agent, creating hybrid DNA origami nanostructures with a stimuli-responsive silica coating for the first time.

In order to test if the properties of the BTDS silicification also hold for large DNA origami superstructures with thicker silica coatings, the octahedral monomers were again polymerised into cubic crystals, followed by BTDS-silicification of the entire crystals. As for the monomers, the crystals were incubated with GSH for 3 h followed by several washing steps, and subsequently subjected to a chelation test. As can be seen from the TEM images in **Figure 6**, the BTDS-silicified crystals exhibit an increased contrast compared to bare crystals, indicating the successful formation of a distinct silica coating. As expected, the BTDS-silicified crystals that were not exposed to GSH also remained intact after the chelation test, successfully demonstrating that the achieved BTDS-silica coating is inferring sufficient stability to the crystals.

**Figure 6:**
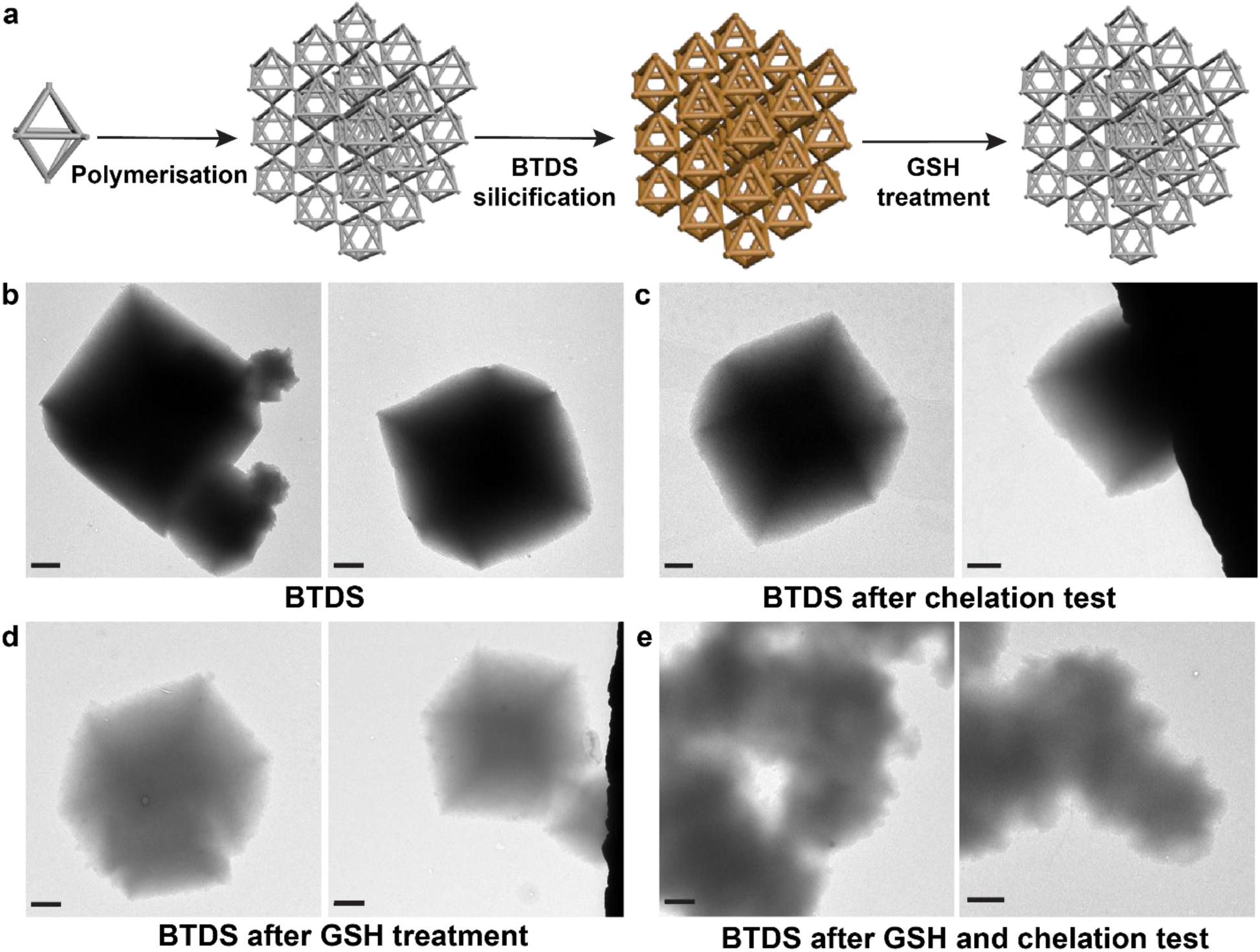
TEM images of BTDS-silicified crystals. (a) Scheme representing the polymerisation of bare octahedral monomers, the subsequent silicification with BTDS and the treatment of the silicified crystals, effectively disrupting the silica coating. (b) crystals directly after silicification with BTDS and removal of excess reagents through washing (c) BTDS-silicified crystals after incubation in a 1x TAE buffer as a chelation test (d) BTDS-silicified crystals after treatment with 10 mM GSH for 3 h. (e) BTDS-silicified crystals after treatment with GSH and subsequent incubation in 1x TAE. While the BTDS-silicified crystals can withstand the chelating conditions, the GSH-treated crystals disassemble during the chelation test, confirming the successful disruption of the BTDS-silica coating by the reducing agent. Scale bars: 1 µm.

Encouragingly, the TEM images of crystals treated with GSH showed a weaker contrast, similar to bare crystals, suggesting the dissolution of the silica shell. This was then further confirmed by the chelation test performed after GSH treatment, displaying the complete disassembly of the crystals in a low-salt environment. These results confirm that BTDS can be successfully used instead of TEOS for the silicification of both DNA origami monomers and large superstructures, providing a protective coating against e.g. low salt environments, but allowing for the dissolution of the silica network in response to treatment with a reducing agent such as GSH.

Again, our findings also shed light on the behaviour of non-standard reagents during silica network formation. In the case of the dissolvable silica, the employed precursor BTDS resembles structurally a TEOS dimer with a connecting disulfide bridge, making the precursor substantially larger in size compared to the TEOS. Similar to the fluorescent silane with the comparably bulky fluorescein moiety, this appears to result in a slower diffusion speed and steric effects negatively affecting the speed of silica growth. Correspondingly, we found that significantly longer incubation times and higher molar excesses of BTDS were required.

Furthermore, the TEM images where the BTDS-silicified crystals appeared less dark than conventionally silicified crystals suggest also a lower shell thickness, despite the higher molar excess of BTDS-silica. The differences in reactivity are also an important consideration when looking at mixtures of different precursors. The observations for crystal silicifications with both TEOS and BTDS in a 1:5 molar ratio indicate that the major part of the silica coating consists of TEOS and the amount of incorporated BTDS is not sufficient to result in a dissolvable shell (**Figure S11c** and **d**).

Thus, we conclude that for the introduction of any silane reagent different from TEOS in a DNA origami silicification, the chemical structure and its potential effects on the reactivity of the reagent and thus on silica network formation have to be taken into account and carefully tuned. Having laid the foundations for such work, we anticipate that this study provides important insights for the future use of other silane precursors.

## Conclusion

In summary, we presented two approaches for the customisation of the silica coating of DNA origami monomers and large crystals. Using two ‘non-standard’ silica precursors that are either commercially available or can be synthesised in a standard chemistry lab, silica coatings of different thicknesses could be endowed with additional functionality: fluorescence and dissolvability.

Firstly, performing a silicification with an additional layer of fluorescent silane on octahedral DNA origami monomers not only increased their stability against heat, chelation and in cell culture medium, but also allowed for the tracking of the silicified structures inside cells. The ability to incorporate fluorescence into the silica coating could thus give additional information about the structural integrity of silicified DNA origami within cells. Moreover, the incorporation of such a protective fluorescent silica coating into DNA origami crystals not only allowed us to study the distribution of the fluorescence in thick silica coatings, but further offers an interesting property for materials science applications.

DNA origami crystals generally provide an interesting platform for many different potential applications, particularly in combination with techniques for the inorganic coating of these structures, as properties such as the lattice type, the solvent channel size or the average size of the crystal grains can be finely tuned.

Secondly, the silica precursor BTDS was employed to create silica coatings that can be controllably dissolved upon exposure to a reducing agent. This was found to be possible even for thick silica coatings on DNA origami crystals. The presented results show that BTDS is an interesting candidate for potential applications e.g. in the field of drug delivery.

Additionally, we anticipate that our study has paved the way for future advances in the customisable silica coating of DNA origami nanostructures. The results here allowed for insights into the behaviour of ‘non-standard’ silica reagents, particularly in terms of silica network formation, and into the required adjustments to experimental protocols. We envision that the palette of additional functionalities for silica coatings of DNA origami can be further expanded using the strategies presented in this study. While fluorescein is an attractive choice, other dyes with different properties such as e.g. rhodamine could be preferable depending on the application. Moreover, other types of stimuli-responsive silica coatings for DNA origami are conceivable that show e.g. light-triggered dissolvability.

## Acknowledgements

The authors thank Lea Wassermann for help with initial experiments and Marianne Braun for assistance with TEM imaging and the preparation of crystal slices. Furthermore, the authors acknowledge the support from Martin Spitaler and the Imaging Facility (RRID:SCR_025739) of the MPI of Biochemistry.

A.H.-J. acknowledges funding from the Deutsche Forschungsgesellschaft (DFG) through the Emmy Noether program (project 427981116) and the SFB1032 “Nanoagents for the spatiotemporal control of cellular functions” (A06). A.B. and L.G. acknowledge support from the IMPRS-ML graduate school. P.M. acknowledges support from the Joachim Herz Foundation (Research Fellowship) and the Studienstiftung des Deutschen Volkes (Ph.D. scholarship). O.T.-S. thanks the German Research Foundation (DFG) for position support through an Emmy Noether grant (project 400324123).

## Supplementary Information

### Note S1: Experimental Methods

#### Synthesis of fluorescent silica precursor

##### Analytical Methods

High resolution mass spectrometry (**HRMS**) was conducted using a *Thermo Finnigan LTQ FT Ultra FourierTransform* ion cyclotron resonance spectrometer from *ThermoFisher Scientific GmbH* applying electron spray ionisation (ESI) with a spray capillary voltage of 4 kV at temperature 250 °C with a method dependent range from 50 to 2000 u.

Nuclear magnetic resonance (**NMR**) spectroscopy was performed using a *Bruker Avance III HD Biospin* (400/100 MHz, with BBFO cryoprobe^TM^) from Bruker Corp. at 400 MHz. NMR-spectra were measured at 298 K, and were analysed with the program *MestreNova 14* developed by *MestreLab Ltd.* ^1^H-NMR spectra chemical shifts (δ) in parts per million (ppm) relative to tetramethylsilane (δ = 0 ppm) are reported using the residual protic solvent (CHD_2_OD in CD_3_OD: δ = 3.31 ppm) as an internal reference. For ^13^C-NMR spectra, chemical shifts in ppm relative to tetramethylsilane (δ = 0 ppm) are reported using the central resonance of the solvent signal (CD_3_OD: δ = 49.00 ppm) as an internal reference. For ^1^H-NMR spectra in addition to the chemical shift the following data is reported in parenthesis: multiplicity, coupling constant(s) and number of hydrogen atoms. The abbreviations for multiplicities and related descriptors are s = singlet, d = doublet, t = triplet, q = quartet, or combinations thereof, m = multiplet and br = broad.

Analytical high performance liquid chromatography (**HPLC**) analysis was conducted using an *Agilent 1200 SL* system *Agilent Technologies Corp.,* Santa Clara (USA) equipped with a DAD detector, a *Hypersil Gold* HPLC column from *ThermoFisher Scientific GmbH*, Dreieich (Germany) and consecutive low-resolution mass detection using a LC/MSD IQ mass spectrometer applying ESI from *Agilent Technologies Corp.,* Santa Clara (USA). For both systems mixtures of water (analytical grade, 0.1 % formic acid) and MeCN (analytical grade, 0.1 % formic acid) were used as eluent systems.

**UV-Vis** spectra were recorded on a Cary 60 UV-Vis spectrophotometer from *Agilent Technologies Inc*., Santa Clara (USA) using 1 cm quartz or PMMA cuvettes. The scan rate was set to 600 nm/min and 2.5 nm slit width was used. Unless stated otherwise, the probes and fluorophores were dissolved in PBS (pH = 7.4, 1 % DMSO) at 10 µM concentration.

**Fluorescence spectroscopy** was performed on a Cary Eclipse Fluorescence Spectrometer from *Agilent Technologies Inc*., Santa Clara (USA) using quartz cuvettes. 480 nm light was used for excitation and spectra were recorded from 490−620 nm. The scan rate was set to 120 nm/min and 5 nm slit width was used. Unless stated otherwise, the probes and fluorophores were dissolved in PBS (pH = 7.4, 1 % DMSO) at 10 µM concentration.

All reactions were performed without precautions in regard to potential air- and moisture-sensitivity and were stirred with Teflon-coated magnetic stir bars. For solvent evaporation a *Laborota 400* from Heidolph GmbH, Schwabach (Germany) equipped with a vacuum pump was used. Reactions were monitored by analytical HPLC.

All chemicals, which were obtained from BLDPharm, Sigma-Aldrich, TCI, Alfa Aesar, Acros, abcr or carbolution, were used as received and without purification. Tetrahydrofuran (THF), dichloromethane (DCM) and dimethylformamide (DMF) were provided by Acros and were stored under argon atmosphere and dried over molecular sieves. TLC control, extractions and column chromatography were conducted using distilled, technical grade solvents.

### Synthesis procedure

Two regioisomers were used in this paper, synthesised by two synthetic procedures: (1) Amide coupling in DMSO which can be directly used for silicification, and (2) Amide coupling in DMF followed by extraction to remove the *N*- hydroxy succinimide side product. Generally, we recommend using **Procedure 1** as it avoids silicification during the product isolation.

### Synthesis fluorescein-5-silane (S01)

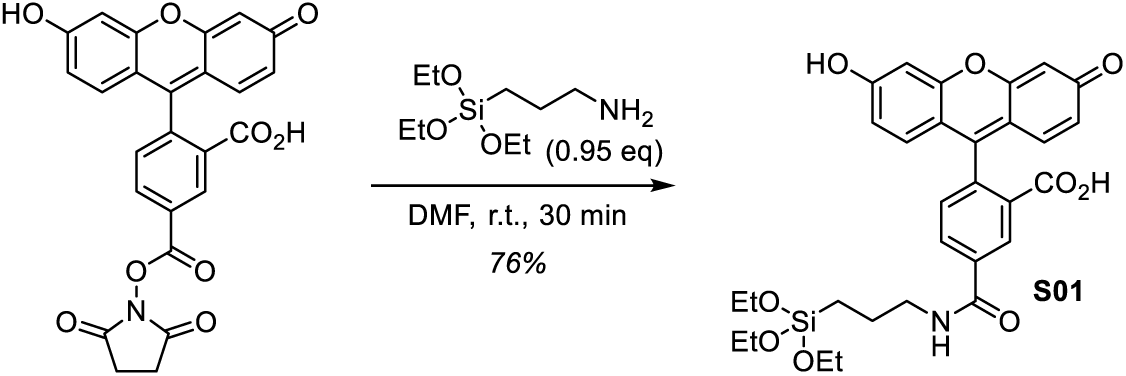

(3-Aminopropyl)triethoxysilane (7.1 µL, 30 µmol, 1.0 equiv.) was diluted with anhydrous, perdeuterated dimethyl sulfoxide (DMSO-d6, 93 µL) and added to 5-carboxyfluorescein *N*-succinimidyl ester (purchased from TCI, 15 mg, 32 µmol, 1.05 equiv.). The mixture was heated to 37 °C in a heat bath for 5 min and then incubated at room temperature for 2 h (without stirring). After full conversion, DMSO-d6 (500 µL) was added to obtain a 50 mM stock solution of **S01**.

**^1^H-NMR** (400 MHz, DMSO-d6): *δ* (ppm) = 8.83 (t, *J* = 5.6 Hz, 1H), 8.52 – 8.42 (m, 1H), 8.24 (dd, *J* = 8.1, 1.5 Hz, 1H), 7.37 (d, *J* = 8.1 Hz, 1H), 6.69 (d, *J* = 2.2 Hz, 2H), 6.59 (d, *J* = 8.7 Hz, 2H), 6.55 (dd, *J* = 8.7, 2.2 Hz, 2H), 3.76 (q, *J* = 7.0 Hz, 6H), 3.29 (q, *J* = 6.8 Hz, 2H), 1.61 (p, *J* = 7.9 Hz, 2H), 1.15 (t, *J* = 7.0 Hz, 9H), 0.65 – 0.58 (m, 2H); **^13^C-NMR** (101 MHz, DMSO-d6): *δ* (ppm) = 168.3, 164.5, 159.7, 154.6, 151.9, 136.4, 134.7, 129.2, 126.5, 124.3, 123.3, 112.7, 109.1, 102.3, 83.4, 57.8, 42.2, 22.6, 18.3, 7.5.

### Synthesis of fluorescein-6-silane (S02)

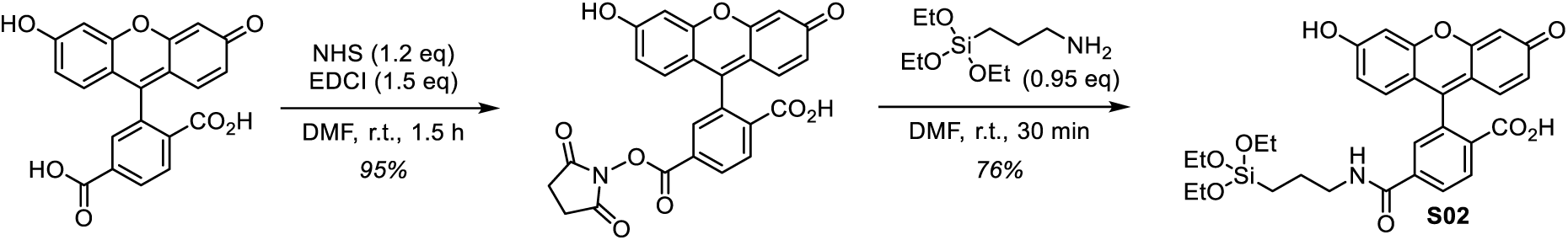

6-carboxyfluorescein NHS-ester was prepared following a previously described procedure.^1^ 6-Carboxyfluorescein (purchased from BLDPharm, 30 mg, 80 µmol, 1.0 equiv.) was dissolved in anhydrous *N*,*N*-dimethyl formamide (2 mL). 1-Ethyl-3-(3-dimethylaminopropyl)carbodiimide hydrochloride (EDCI, 23 mg, 0.12 mmol, 1.5 equiv.) and *N*- hydroxysuccinimide (11 mg, 96 µmol, 1.2 equiv.) were added and the reaction was stirred at room temperature for 1.5 h. The reaction mixture was diluted with ethyl acetate (20 mL) and washed with phosphate buffer (pH = 6, 2 × 20 mL). The organic layer was dried over sodium sulfate, the desiccant was filtered off and the volatiles were removed *in vacuo* to give 6-carboxyfluorescein NHS-ester (36 mg, 76 µmol, 95%) as an orange solid.

The obtained 6-carboxyfluorescein NHS-ester was dissolved in anhydrous *N*,*N*-dimethyl formamide (2 mL). (3-Aminopropyl)triethoxysilane (17 µL, 71 µmol, 0.95 equiv.) was added and the reaction mixture was stirred at room temperature for 30 min. The mixture was diluted with ethyl acetate (30 mL) and water (30 mL) and the layers were separated. The organic layer was dried over sodium sulfate, the desiccant was filtered off and the volatiles were removed *in vacuo* to give the title compound (**S01**, 30 mg, 52 µmol, 76%) as an orange solid.

*<u>Note</u>: the reaction was monitored by HPLC analysis regarding consumption of the NHS-ester. The product however decomposes during the HPLC run to partially reveal the de-ethylated R-Si(OH)_3_ decomposition product. This does not correspond to decomposition during the reaction as confirmed by NMR analysis of the product*.

**^1^H-NMR** (400 MHz, MeOD): *δ* (ppm) = 8.13 (dd, J = 8.0, 1.4 Hz, 1H), 8.09 – 8.06 (m, 1H), 7.60 (s, 1H), 6.69 (d, J = 2.3 Hz, 2H), 6.61 (d, J = 8.7 Hz, 2H), 6.54 (dd, J = 8.7, 2.4 Hz, 2H), 3.75 (q, J = 7.0 Hz, 6H), 3.31 – 3.24 (m, 2H), 1.69 – 1.58 (m, 2H), 1.15 (t, J = 7.0 Hz, 9H), 0.65 – 0.56 (m, 2H); **^13^C-NMR** (101 MHz, MeOD): *δ* (ppm) = 170.6, 168.2, 161.4, 155.0, 154.1, 142.5, 130.3, 130.3, 130.3, 126.2, 123.9, 113.8, 110.9, 103.6, 85.7, 59.5, 43.7, 23.7, 18.6, 8.6; **HRMS** (ESI−): m/z calc. for C_30_H_32_NO_9_Si^−^ [M−H]^−^: 578.1852, found: 578.1851.

### Folding and purification of DNA origami octahedra

The DNA origami octahedra used here were designed using the TALOS^2^ and CaDNAno^3^ softwares (design schematics in **Figure S2**).

The folding of the octahedra was carried out using 20 nM of the scaffold p7249 and 100 nM of each staple strand in buffer containing 40 mM Tris, 20 mM acetic acid, 1 mM EDTA (pH = 8.0) and 15 mM MgCl_2_. Two separate mixtures were prepared, containing the two different sets of end staples (type A or type B). The mixtures were heated to 95°C and held at this temperature for 1 min and then cooled down to 20 °C over a period of 20 h. The fully assembled DNA origami nanostructures were purified from excess staple strands via ultrafiltration (Amicon centrifugal filter units, 0.5 ml, 100 kDa cut-off). The folding mixtures were loaded into pre-wetted filter units (type A and B separately, 250 µL per filter unit together with 250 µL of fresh buffer) and centrifuged at 2,000 rcf for 20 min. The centrifugal steps were repeated 5 times, whereby fresh buffer (1 x TAE, 7.5 mM MgCl_2_, 400 µL) was added in every step. Finally, the DNA origami were eluted by placing the inverted filter unit into a new collection tube and spinning the tube for 8 min at 2,000 rcf.

The successful folding of the octahedral nanostructures was confirmed by TEM imaging.

### Polymerization of octahedral DNA origami monomers / DNA origami crystal formation

Assembly of cubic crystals from octahedral DNA origami monomers was achieved by mixing the two different monomer types (A and B, at least 15 nM each, either both bare or both silicified) in a 1 x TAE buffer containing 25 mM MgCl_2_. The reaction mixture (total volume 40 µl) was heated to 48 °C for 1 h and then slowly and gradually cooled down to 20 °C over a period of around 70 h (-1 °C per 150 min).

### Silicification of DNA origami octahedra (monomers)

The silicification of octahedral DNA origami monomers was performed similarly to the procedure described previously.^4^ 50 µL of purified DNA origami sample were prepared at a concentration of 50 nM in buffer containing 1 x TAE, 7.5 mM MgCl_2_ in a round-bottom Eppendorf tube. The tube was placed on a thermo shaker and the first silica precursor TMAPS (TCI, diluted 1:19 in methanol) was added to the solution in a 6-fold molar excess to the number of nucleobases. After one minute of shaking at 400 rpm at 21 °C, the second silica precursor TEOS (Sigma Aldrich, diluted 1:9 in methanol) was added to the sample in a 15-fold molar excess to the number of nucleobases. The reaction mixture was left on the thermo shaker for 15 min at 400 rpm and was then transferred to a V-bottom tube and put on a tube revolver rotator (Thermo Scientific) and rotated at 40 rpm at room temperature (21°C) for 6 h.

Subsequently, the sample was purified from excess silica via buffer exchange in spin filtration columns (Zeba Spin Desalting Columns, 0.5 mL, 40 kDa cut-off). After the storage solution was eluted from the spin columns (1,500 rcf, 1- 2 min), they were washed three times with fresh buffer (1 x TAE, 7.5 mM MgCl_2_). Then, the spin columns were placed in a fresh collection tube and the sample was added into the columns. After centrifugation at 1,500 rcf for 2 min, the eluted sample was collected.

For fluorescent silicification of the DNA origami octahedra, the silicified (and purified) sample (ca. 50 µL, 45 nM) was placed into a light-protected V-bottom tube and 7.5 µL of fluorescent silica precursor (10 mM) were added. After careful mixing of the solution, the tube was again placed on the tube revolver rotator and rotated at 40 rpm for the desired incubation time (3 d). Purification was carried out using up to two rounds of spin filtration, as described above.

For the BTDS-silicification of the octahedral monomers, the BTDS was added undiluted to the solution on the thermo shaker in a 125-fold molar excess to the number of nucleobases instead of the TEOS. Furthermore, the incubation time on the tube revolver rotator was extended to 4 d.

### Silicification of DNA origami crystals made from octahedral monomers

A sample with freshly polymerized DNA origami crystals (40 µl, 30 nM monomers) was put into a round-bottom tube and the tube was placed on a thermo shaker at 4 °C and shaken at 700 rpm. 0.75 µl of the first silica precursor TMAPS (diluted 1:9 in methanol) were added to the solution and the sample was left shaking for 30 min. Then, 0.75 µl of the second silica precursor TEOS were added to the sample and the shaking was continued for another 210 min. Then, the temperature was increased to 10 °C, while the shaking speed was kept at 700 rpm for another 2 h. After that, the temperature was further raised to 16 °C and the shaking was continued for 2 h. Following that, the temperature was increased to room temperature (21 °C) and the shaking speed was reduced to 300 rpm. Then, the sample was left shaking overnight (16 h, total time 24 h).

For the purification of the silicified crystals, the sample was transferred to a V-bottom tube and 300 µl of buffer solution were added. Then, the sample was left undisturbed, so that the silicified can sediment at the bottom of the tube. In one washing step, the supernatant (ca. 300 µl) was carefully removed from the top of the solution, followed by a refilling of the buffer solution. For each sample, at least 3 washing steps were carried out.

For fluorescent silicification of the crystals, 1 – 7.5 µl of fluorescent silica precursor (10 mM) were added during the shaking (see **Supplementary Figure S5**, optimal time point: 10 h after start of the silicification procedure).

Purification was carried out by transferring the sample into a new V-bottom tube and adding 300 – 500 µl of fresh buffer (1 x TAE, 7.5 mM MgCl_2_). The sample was left undisturbed until the silicified crystals sedimented at the bottom of the tube. Then, the liquid above the crystals was very carefully removed and the tube was re-filled with fresh buffer. This washing step was repeated at least twice. All samples containing Cy5 and / or fluorescent silica were kept in the dark / in light-protected tubes.

For the fabrication of a dissolvable silica coating, the TEOS was replaced by 1.86 µl of undiluted BTDS that was added undiluted after 30 min of incubation on the thermo shaker.

### Agarose Gel Electrophoresis

DNA origami samples (15 µl, 10 nM in 1 x TAE buffer with 7.5 mM MgCl_2_) were mixed with loading buffer containing Orange G and Ficoll and loaded onto a 0.7% agarose gel (1 x TAE, 11 mM MgCl_2_). The gel was run on ice for 120 min at 75 V (running buffer: 1 x TAE with 11 mM MgCl_2_). Gel imaging was subsequently carried out using the Typhoon FLA-9000 (GE Healthcare). For post-staining, the gel was incubated in 100 mL MilliQ water with 0.01 % SYBR Gold for 20 min and then thoroughly washed in MilliQ water, followed by another round of imaging.

### Transmission Electron Microscopy

Monomers: A DNA origami sample (10 µl, 10 nM) was applied to a carbon-coated copper grid (300-mesh, Plano GmbH) that had been plasma-cleaned for 30 s. Silicified DNA origami samples were incubated on the TEM grids for 6- 8 min. In the next step, the remaining solution was carefully removed from the grid using filter paper and the grid was washed with MilliQ water (5 µl each) twice and dried in air before imaging.

Crystals: The DNA origami crystals were collected from the bottom of a tube and the sample (10 µL) was then incubated on a plasma-cleaned (30 s) TEM grid for 45 to 55 min. Subsequently, the remaining liquid was carefully removed with filter paper and the grid was washed twice with MilliQ water (5 µl each) and then air-dried before imaging.

All images were recorded on a Jeol-JEM-1230 TEM operating at an accelerating voltage of 80 kV with a Gatan Orius SC1000 digital camera (software: Gatan DigitalMicrograph). The images were subsequently analyzed using the ImageJ software^5^.

Preparation of crystal slices: The silicified DNA origami crystals were collected from the bottom of a tube (10 µl) and placed into a fresh V-bottom tube. A solution containing 4% agarose was prepared and kept on a heat plate for a few minutes to avoid solidification and remove as much air as possible from the solution. Subsequently, approximately 100 µl of the agarose solution were carefully poured into the tube that contained the sample in order to encapsulate the DNA origami crystals in the agarose. Then, the sample was left at room temperature for full solidification. In the next step, small cubes (2 mm x 2 mm x 2 mm) containing the crystals were cut out. These cubes were then successively immersed in increasing amounts of ethanol (30%, 50%, 70%, 96%, 2x 100%) for 20 min per step, followed by the embedding in Spurr’s low viscosity embedding medium according to the manual. After the embedding, the crystals were localized in the resin block and a small pyramid (500 µm x 500 µm) was trimmed using a razorblade. Then, slices with a thickness of about 50 nm to 60 nm were cut from these pyramids with a Diatome Diamond Knife (Ultra 45°) using an Ultramicrotome (Leica UC6). The resulting slices were collected on Formvar coated copper slot grids (Plano GmbH).

### Confocal microscopy

The samples for confocal microscopy were prepared in a six-channel ibidi slide. First, 100 µl of BSA-biotin (1 mg/mL, dissolved in buffer containing 10 mM Tris, 100 mM NaCl, pH 8) were incubated in each channel for 10 min, followed by washing with 450 µl of buffer. After that, 100 µl of neutravidin (0.5 mg/mL in buffer containing 10 mM Tris, 100 mM NaCl, pH 8) were added to the channels for 10 min. Following this, the channels were washed with 300 µl of buffer containing 10 mM Tris and 100 mM NaCl and then with 450 µl of buffer containing 1 x TAE and 7.5 mM MgCl_2_. In the next step, the biotinylated DNA origami samples were incubated in the channels for 30 min. Subsequently, the channels were flushed with 450 µl buffer (1 x TAE, 7.5 mM MgCl_2_) and the slide was kept in the dark until the imaging.

The imaging was carried out on a Leica Stellaris 5 confocal microscope with a 63x oil-immersion objective. The obtained images were analyzed with the Fiji Software^6^.

### Cell Culture

HeLa cells (ACC 57, acquired from Leibniz Institute DSMZ) were cultured in DMEM (with high glucose, pyruvate, GlutaMax, added Pencillin and Streptomycin, and 10% FBS) at 37 °C and 5% CO_2_ and reseeded every two to three days with a ratio of 2×10^5^ to 5×10^5^. Cell detachment was conducted using Trypsin/EDTA, during each split cell count and viability were determined using Trypan Blue and an automatic cell counter.

### Uptake experiments

Two days prior to t_0_, 5000 HeLa cells were seeded in 50 µL DMEM (with high glucose, pyruvate, GlutaMax, added Pencillin and Streptomycin, and 10% FBS) in an ibidi 15 well µ-slide and incubated at 37 °C and 5% CO_2_. One day prior to t_0_ the medium was exchanged by FluoroBrite DMEM (supplemented as DMEM). Subsequently, a given volume of medium was removed and exchanged by DNA origami in DPBS with the Mg^2+^ concentration adjusted to 2.5 mM for the 24 h incubated sample. Shorter incubation times were realised by adding the origami suspension at a later time point. The origami concentration in buffer was adjusted to be 30 nM, the exchanged volume was chosen to result in the desired concentration in medium. At t_0_ the cells were washed with cold DPBS (with Ca^+^ and Mg^2+^) and subsequently incubated in Image-iT 4% Paraformaldehyde solution for 15 min at room temperature. Afterwards, the wells were washed with DPBS with 500 ng/mL Hoechst 33342 and again incubated for 5 min at room temperature. After a final wash with DPBS, the cells were imaged using a Leica Stellaris 5 confocal microscope with a 63 × 1.4 oil-immersion lens. Images were deconvoluted using SVI Huygens and further processed in Fiji^6^.

**Figure S1:**
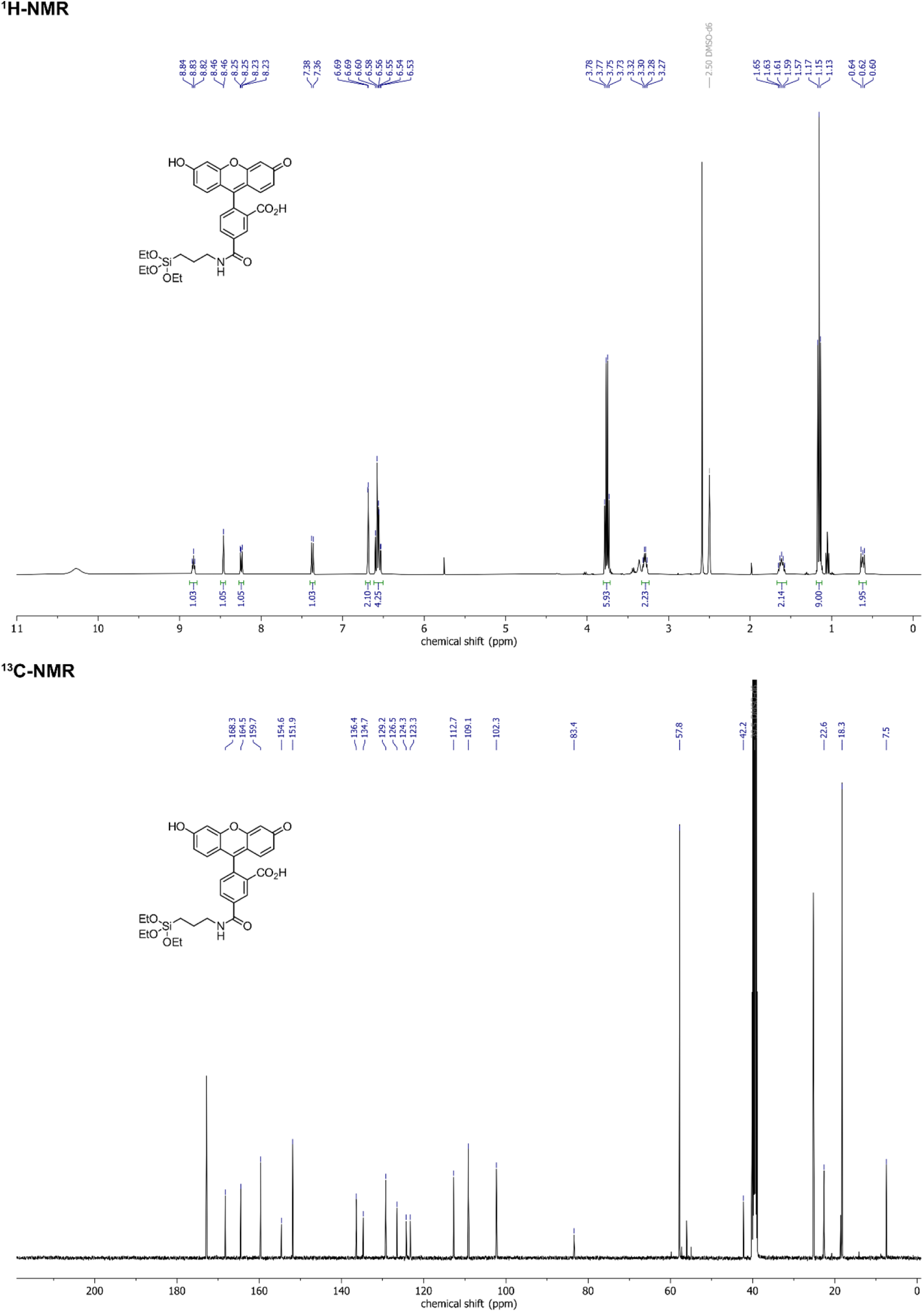
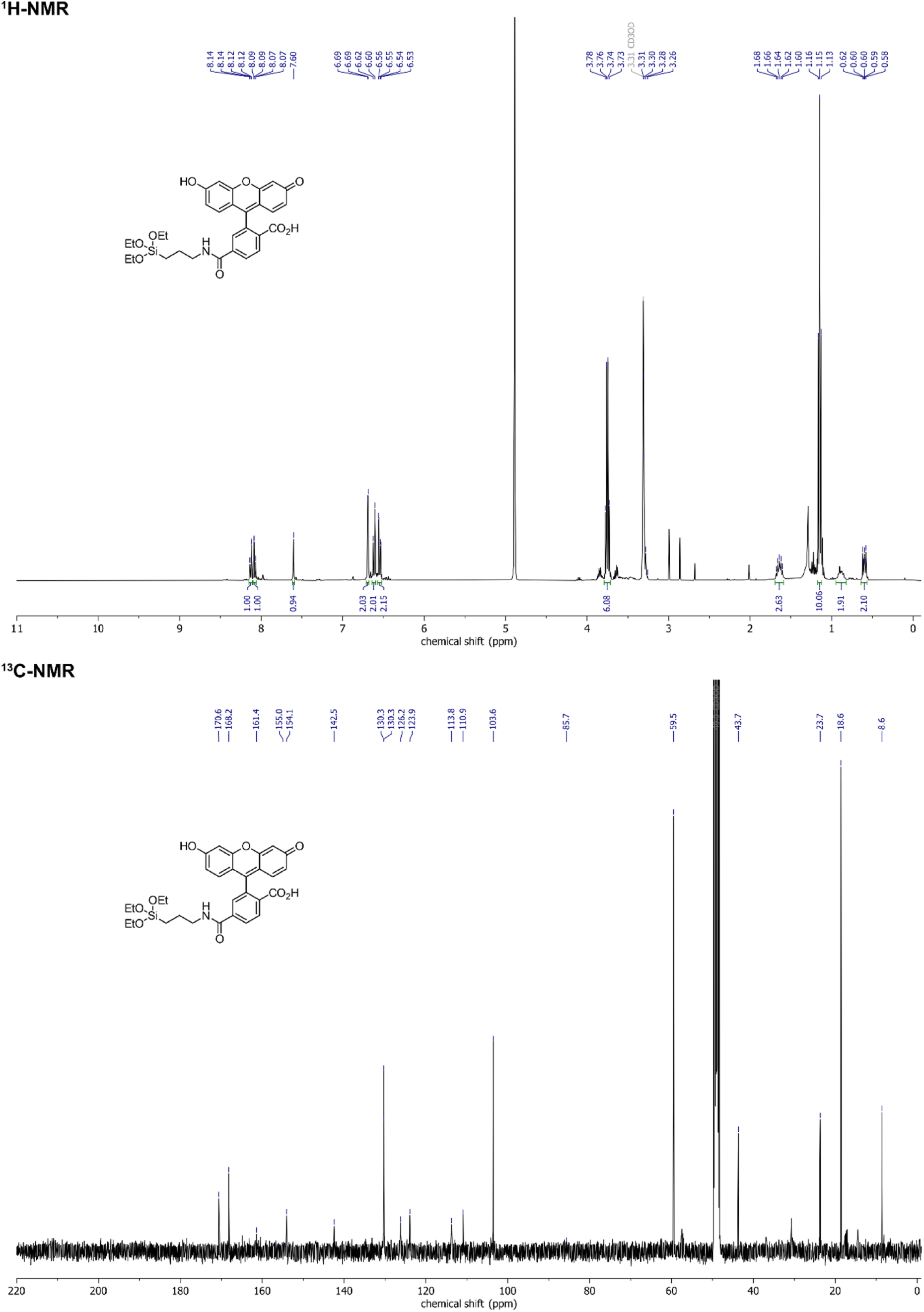
NMR spectra for the two variants of the fluorescent silica precursor (upper images: fluorescein-5-silane (S01), lower images: fluorescein-6-silane (S02)).

**Figure S2:**
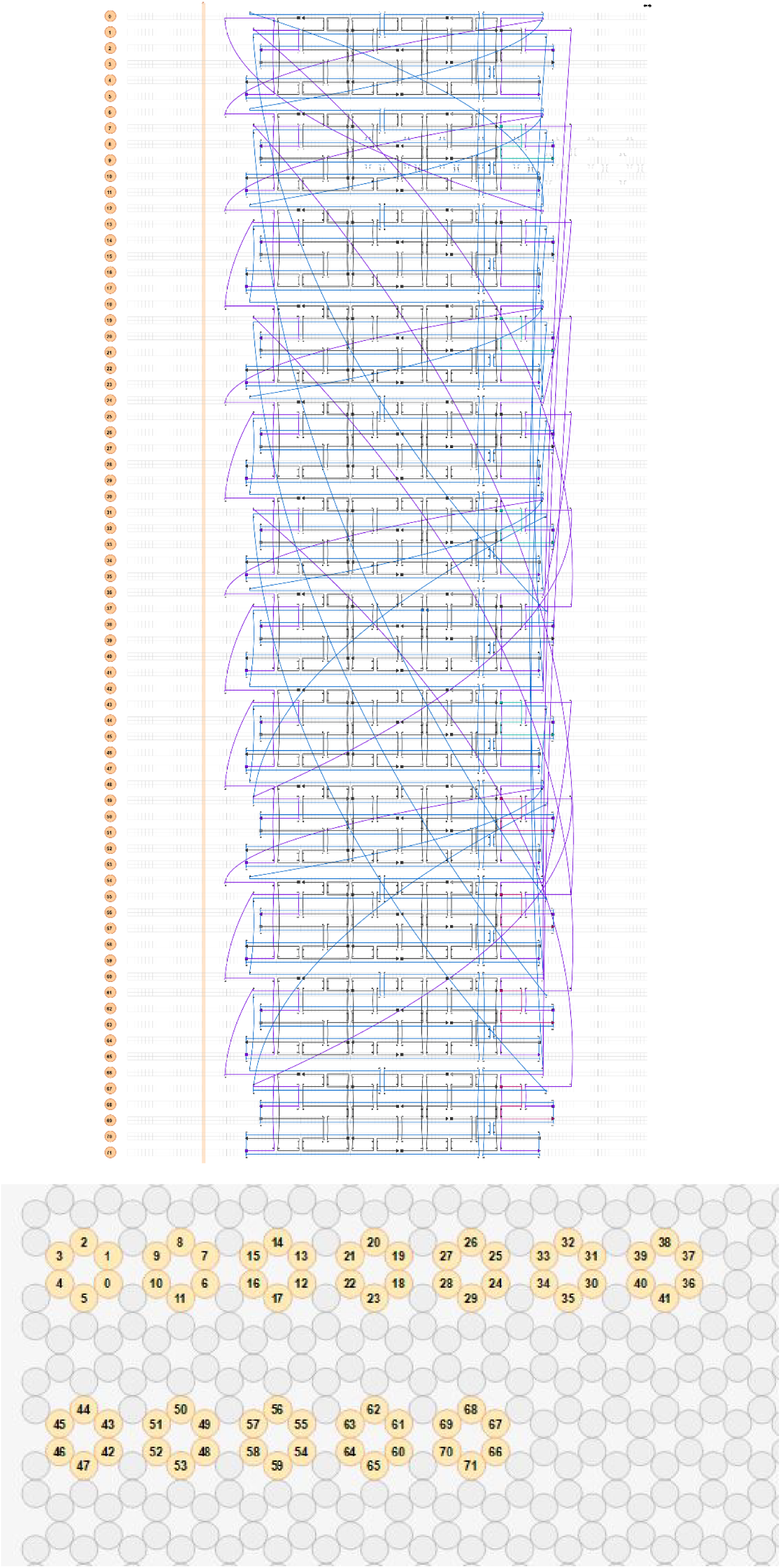
Scaffold routing (blue) and staple strands (black: core staples, purple: sticky ends, magenta: internal Cy5 modifications, light blue: internal FITC modifications) for the DNA origami octahedra consisting of six helix bundles (6HBs). This design was created with the help of the softwares TALOS^2^ and caDNAno^3^. It should be noted that the positioning of the internal fluorescent modifications within the DNA origami could potentially have a significant influence on the timepoint when fluorophores are detached from the nanostructure. Here, the locations of the fluorophore-carrying staple strands at the outer parts of the 6HBs were selected to showcase a rapid loss of signal overlap.

**Figure S3:**
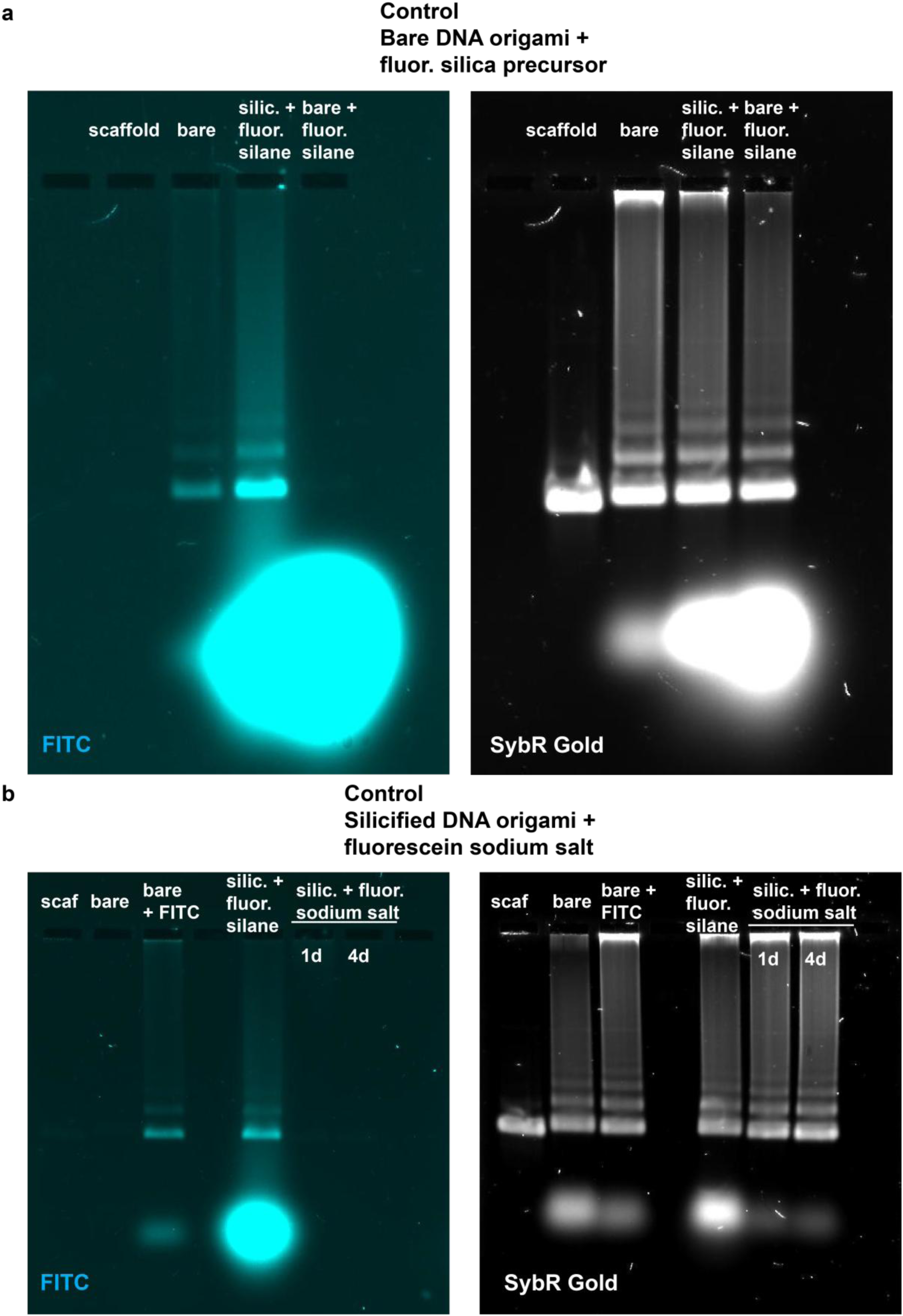
Different control experiments to test for unspecific binding of the fluorescent silane to either the bare DNA origami or of fluorescein itself to the silica shell. (a) AGE showing the effect of the addition of the fluorescent silane to bare DNA origami octahedra with a subsequent overnight incubation (lane on the right). The band for the DNA origami nanostructures does not exhibit a fluorescent signal in the fluorescein channel, confirming that there is no noticeable unspecific binding of the fluorescent silane to the bare DNA origami. (b) AGE for control samples that were silicified (standard silicification) and then incubated for 1 or 4 d, respectively, with fluorescein sodium salt at a similar amount compared to the fluorescent silane. Also here, no fluorescent signal in the lanes for these control samples can be observed, indicating that the fluorescein molecule alone cannot attach to the silica coating of the nanostructures.

**Figure S4:**
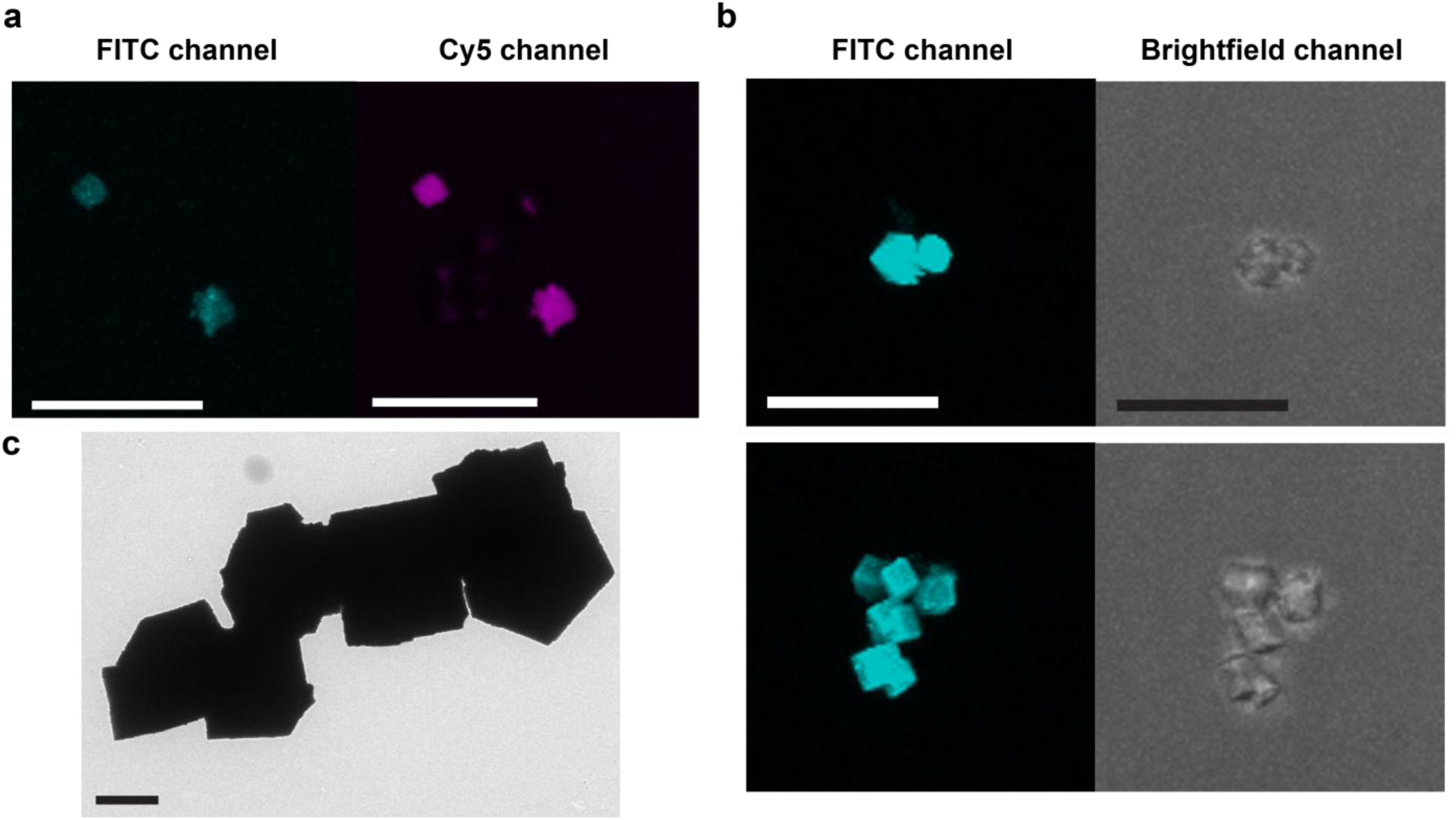
Additional data for fluorescently silicified DNA origami crystals. (a) Confocal microscopy images of crystals assembled from fluorescently silicified octahedral monomers. (b) Confocal microscopy images of crystals that were silicified post-assembly. (c) TEM micrograph of fluorescently silicified crystals with a thick silica coating. Scale bars: 20 µm (a), (b) and 2 µm (c).

**Figure S5:**
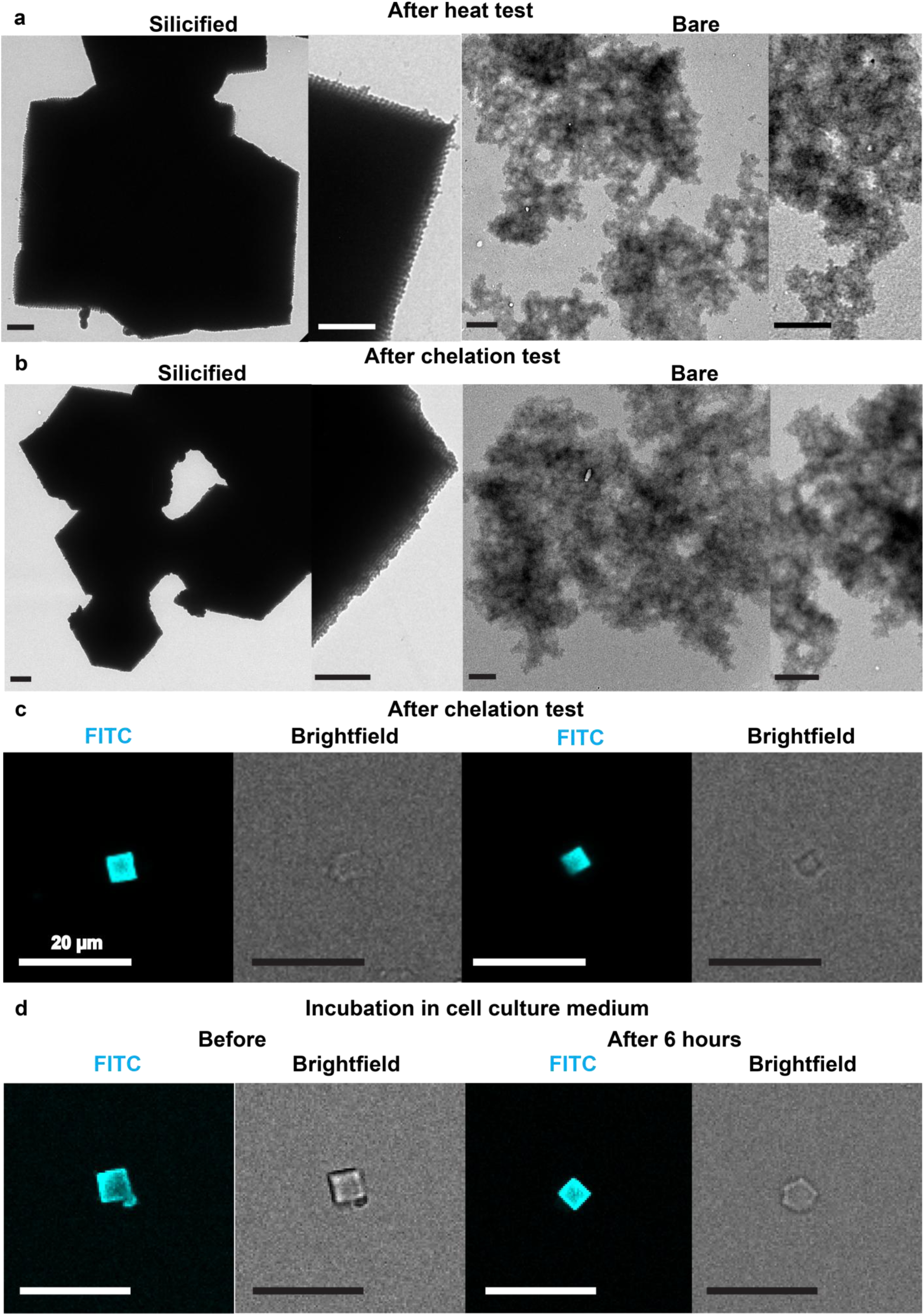
Stability tests for the fluorescently silicified DNA origami crystals. As expected, the fluorescently silicified crystals with their comparatively thick silica deposition can withstand the exposure to elevated temperatures (a, heat test) and low salt conditions (b, chelation test), while bare crystals fall apart. The stability of the crystals upon incubation in 1x TAE buffer (c) and DMEM with 10% FBS (d) was further monitored with the help of confocal microscopy. Even after several hours of exposure to detrimental environmental cues, the crystals still exhibit a strong fluorescent signal in the fluorescein channel. Scale bars: 1 µm (a), (b) and 20 µm (c), (d).

**Figure S6:**
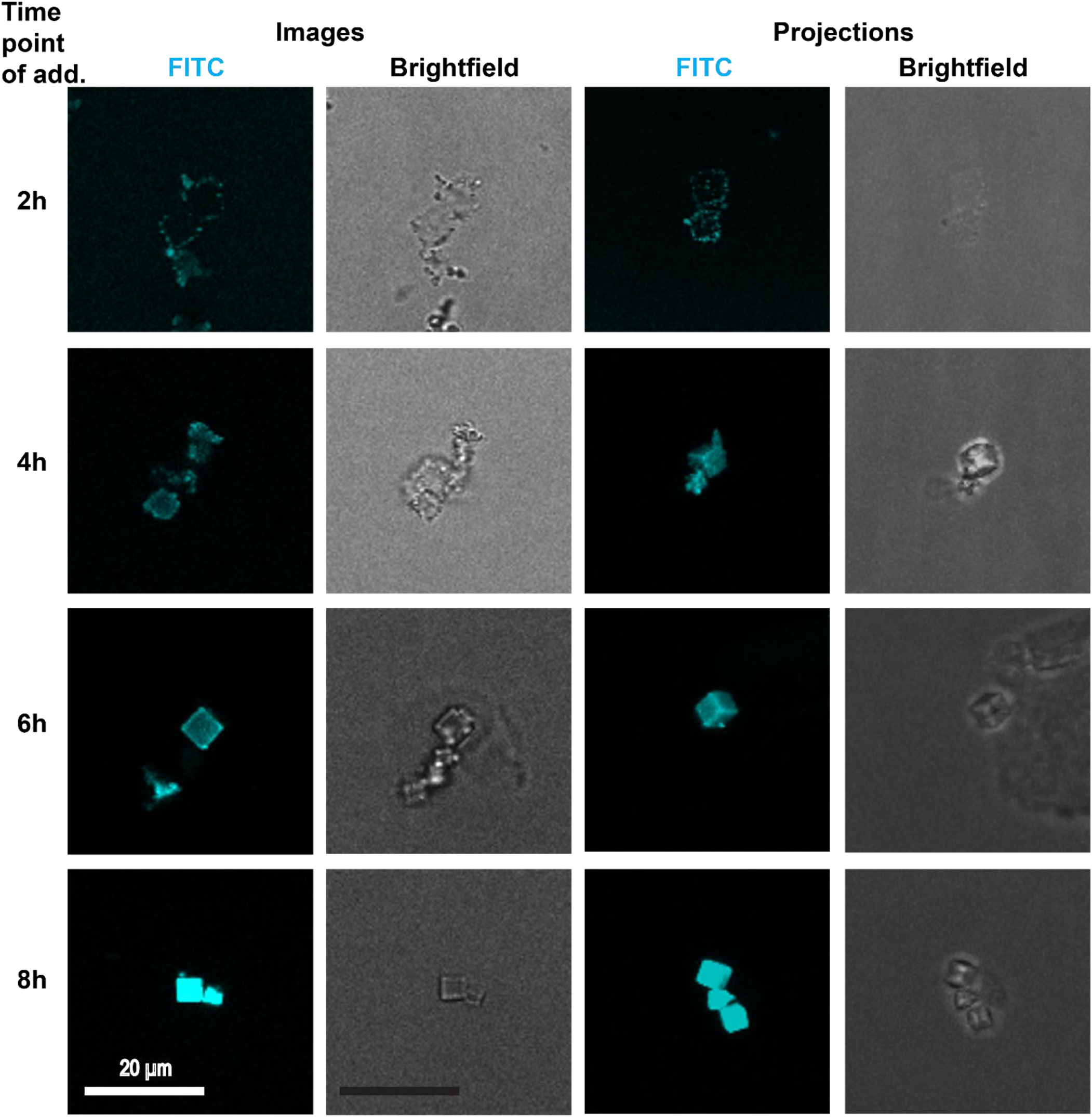
Confocal microscopy images for silicified crystals assembled from octahedral DNA origami monomers with different time points of fluorescent silane addition. Left columns: single images, right columns: 3D projections of z-stacks. When the fluorescent silane is added only 2 h after the start of the silicification, the fluorescent silica does not reach the centre of the cubic crystals, but appears to be forming silica clusters on the outside of the structures. For the fluorescent silica addition after 4 h, there is still the formation of silica clusters around the crystals visible, even though there is also a (weak) fluorescent signal from the crystal centres. More homogeneous results can be achieved with an even later addition of the fluorescein-modified reagent, as it can be seen from the results in two lower rows. Scale bars: 20 µm.

**Figure S7:**
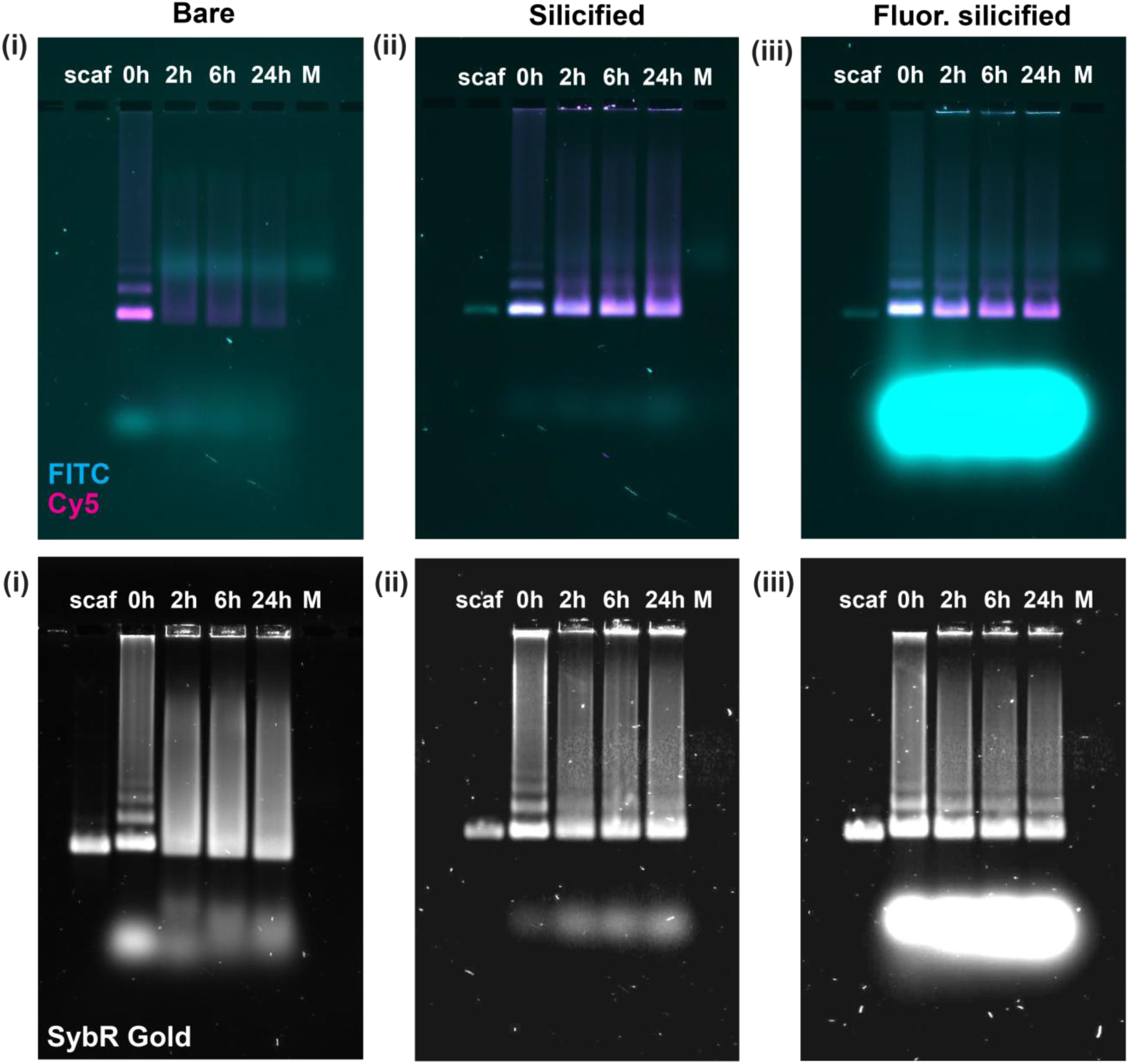
AGE for bare, silicified and fluorescently silicified DNA origami samples after the incubation in DMEM with a final concentration of 10% FBS for up to 24 h. (i) The bare octahedron with both Cy5 and FITC internal modifications show clear signs of disassembly. Both the Cy5 and the FITC signal decrease rapidly (within the first 2 h) and a band that can probably attributed to the degradation of octahedron into 6HBs appears above the staple pools. (ii) In the silicified sample, there is still colocalization of the signals from the internal fluorophores even after 24 h and there are no additional bands appearing below the DNA origami bands, showing that the silica shell significantly enhances the stability of the nanostructures. (iii) Also the fluorescently silicified octahedra appear to be stable and show a substantial overlap of the Cy5 and the fluorescein signal from the silica shell after 24 h in the cell culture medium. However, the fluorescein signal is decreasing over time, indicating that still some part of the originally bound fluorescent silane is detached from the nanostructures with increasing incubation time.

**Figure S8:**
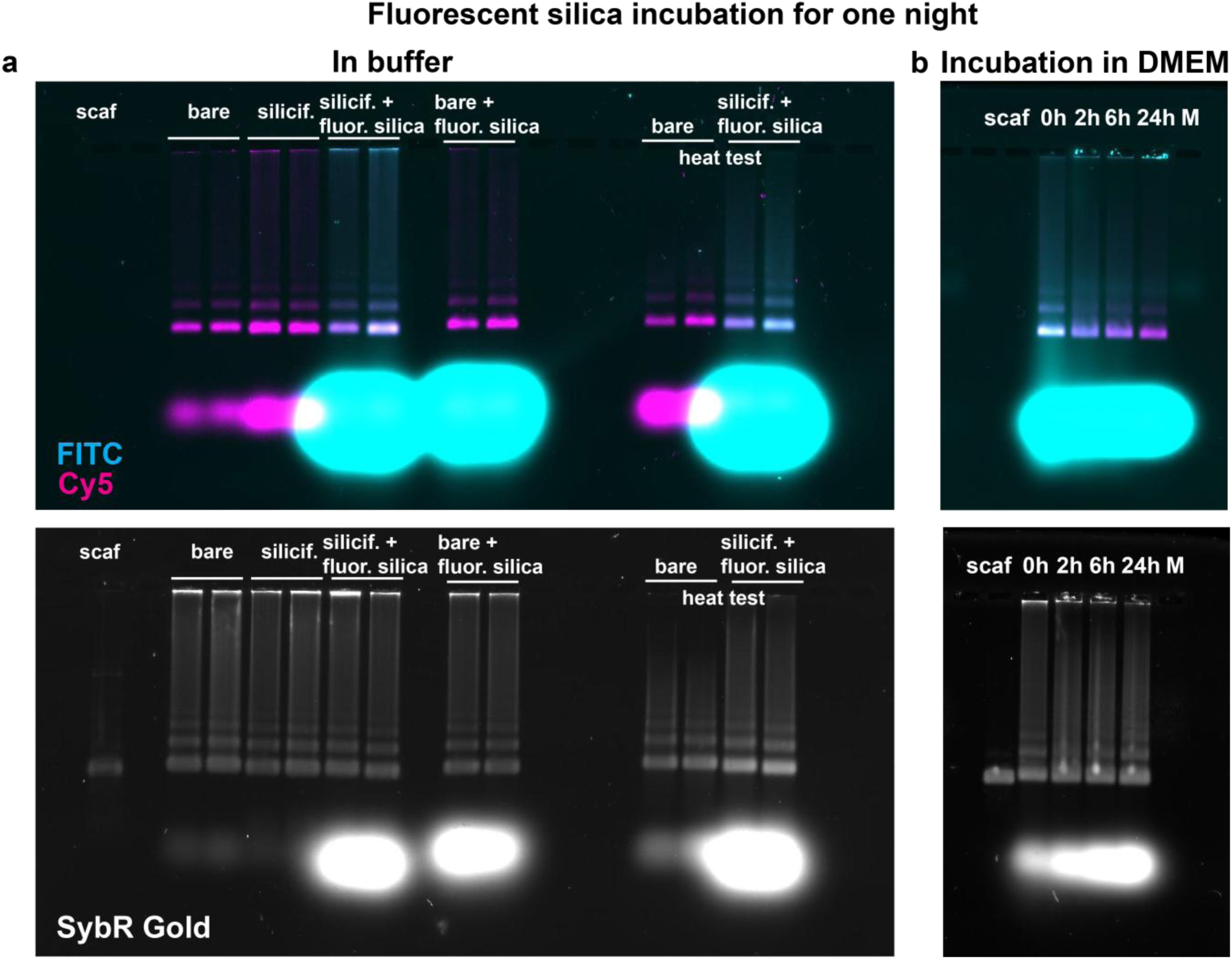
Behaviour of silicified DNA origami octahedra after only one overnight incubation with the fluorescent silane. (a) AGE showing the strong fluorescent signal in the DNA origami band for the fluorescently silicified octahedra. The control samples, where only methanol instead of TMAPS and TEOS was added during the first silicification round, do not exhibit a signal in the fluorescein channel and behave similarly to the bare and silicified (TEOS only) samples with internal Cy5 modifications. The fluorescently silicified octahedra exhibit a good stability upon exposure to heat (60 °C for 30 min). (b) AGE showing fluorescently silicified DNA origami samples after incubation in DMEM with a final concentration of 10% FBS for up to 24 h. Strikingly, the signal in the fluorescein channel decreases strongly over time while the Cy5 signal stays in a similar range, suggesting that the outer fluorescent silica layer is lost during incubation in the cell culture medium.

**Figure S9:**
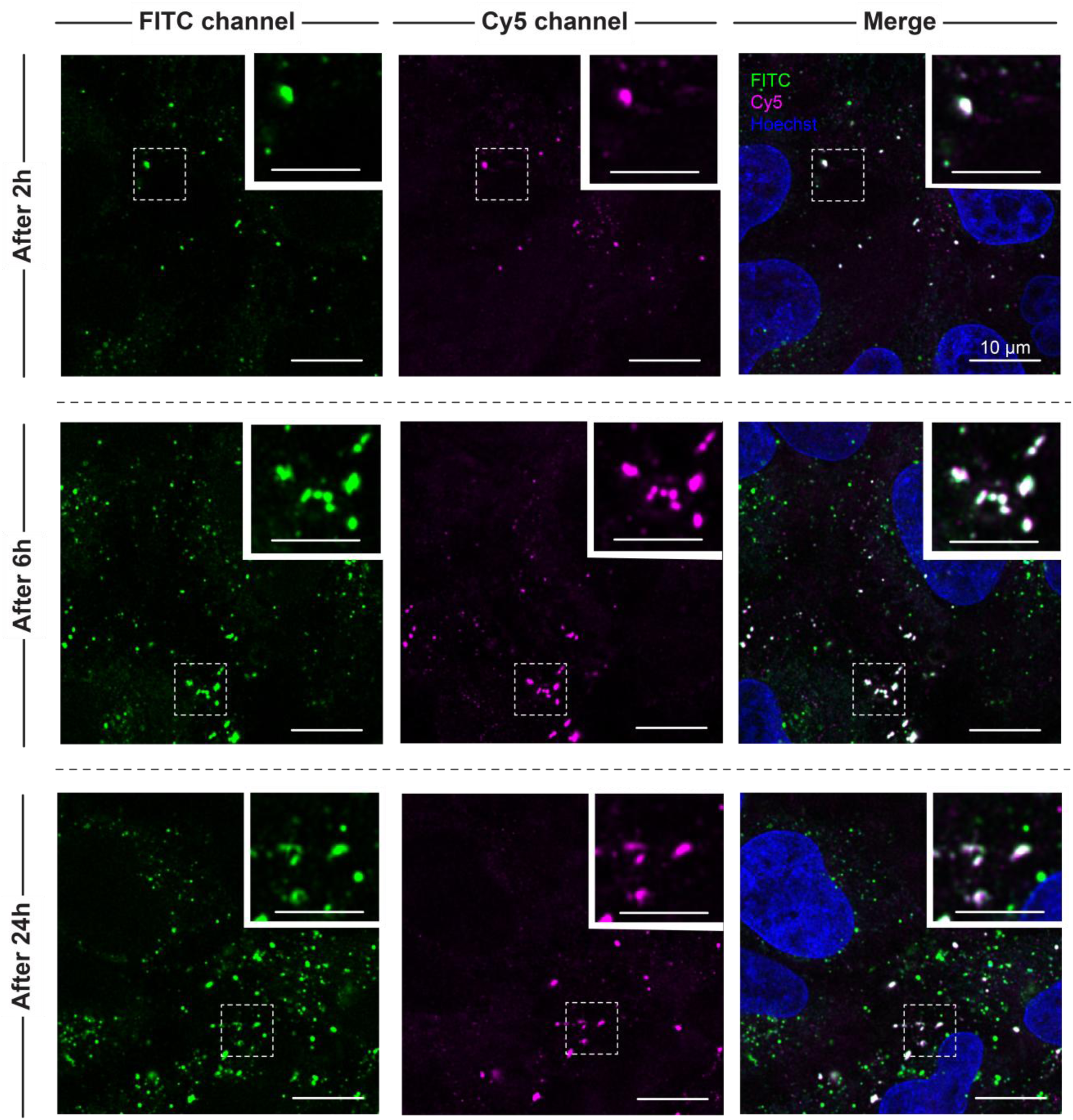
Additional data for the cellular uptake of fluorescently silicified DNA origami octahedra after 2 h, 6 h and 24 h. Scale bars: 10 µm, insets: 5 µm.

**Figure S10:**
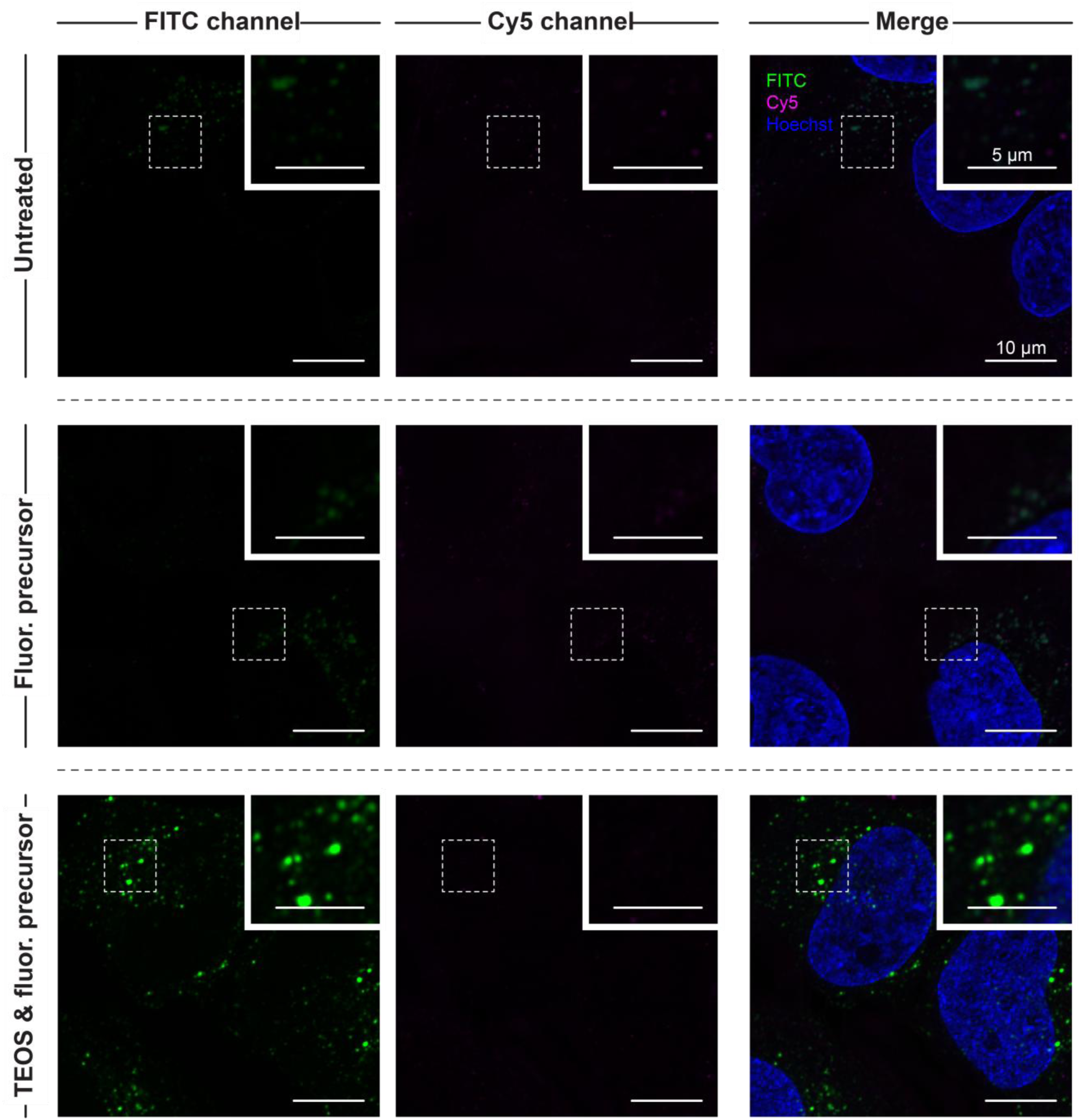
Control experiments for the uptake of the fluorescent silica precursor without DNA origami. The control sample with only buffer (upper row) and the sample with fluorescent silane taken directly from the stock solution (middle row) do not exhibit a significant fluorescent signal. Contrary to that, the sample containing TMAPS, TEOS and fluorescent silane after incubation in buffer for 4 d and subsequent purification (lower row) shows bright fluorescent spots, confirming that silica clusters with fluorescent silica can form during the silicification procedure and are not entirely removed during the purification steps. Scale bars: 10 µm, insets: 5 µm.

**Figure S11:**
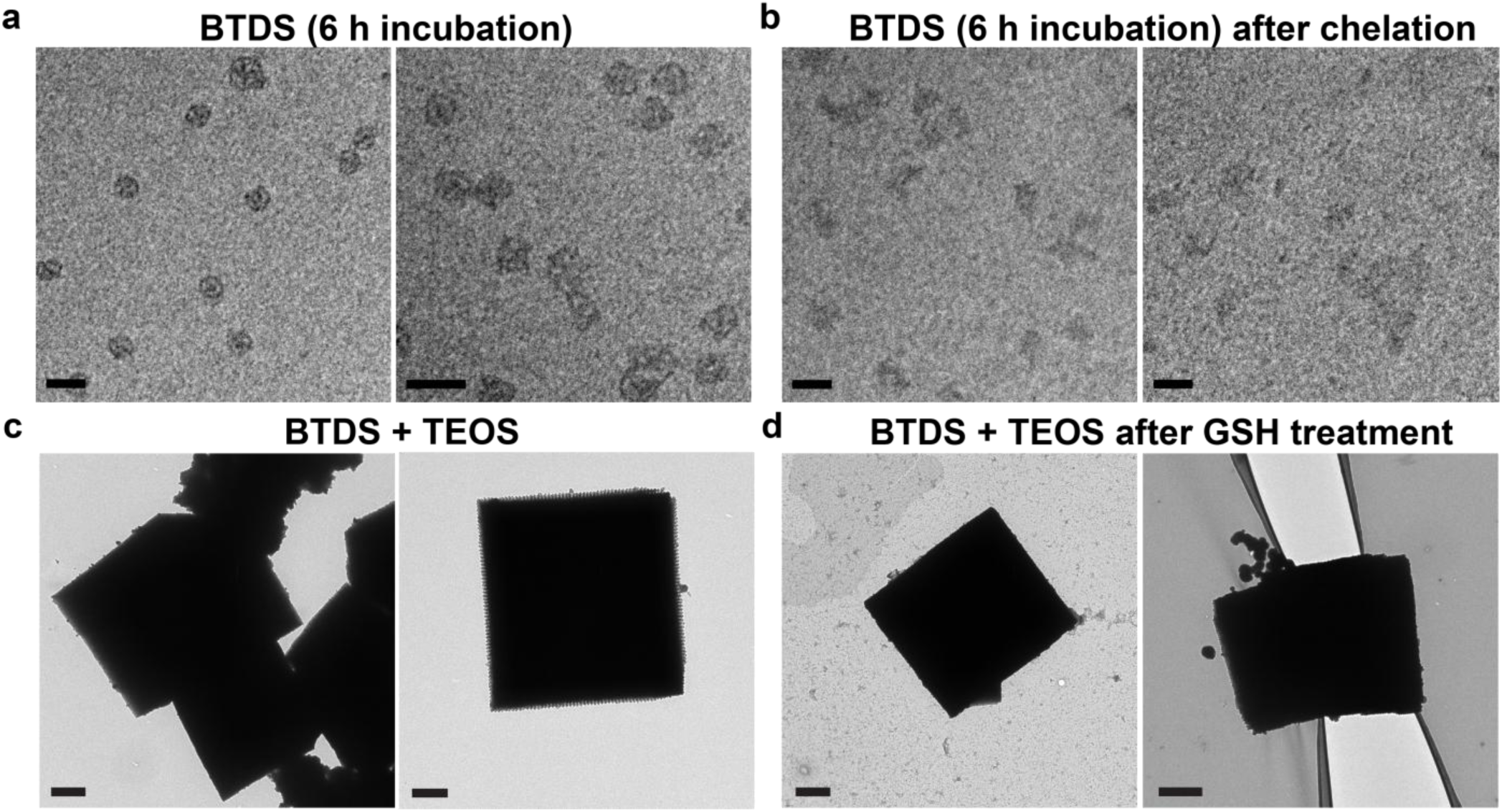
(a) Octahedral DNA origami monomers after a ’standard’ silicification procedure with BTDS instead of TEOS and only 6 h of incubation with the silica reagents. The resulting structures do not exhibit sufficient stability against chelation (b), indicating the need for a prolonged duration of the silicification procedure. Scale bars: 100 nm. (c) TEM micrographs showing cubic crystals that were silicified post-assembly with a mixture of BTDS and TEOS in a 5:1 ratio. The crystals appear to have a thick silica coating. (d) TEM images of such crystals after incubation with glutathione. The silica coating does not exhibit any signs of disruption or dissolution, suggesting that the silica layer consists predominantly of TEOS that resists the GSH treatment. Scale bars: 1 µm.

**Figure S12:**
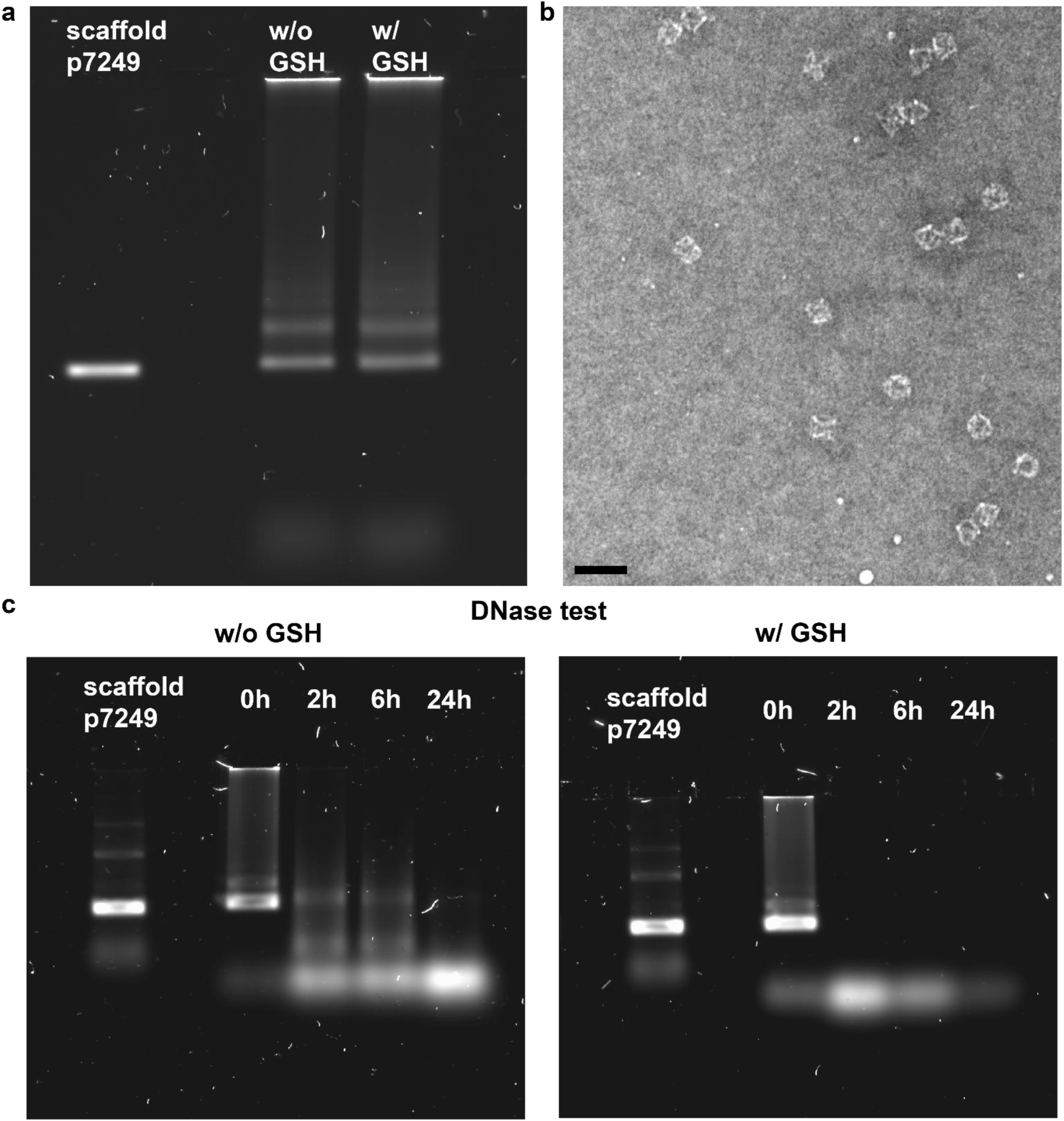
Effect of GSH on octahedral DNA origami monomers. (a) Agarose gel showing bare octahedra that were incubated in 10 mM GSH for 3 h, followed by a purification step. There is no noticeable difference between the untreated and the GSH-treated DNA origami in the agarose gel. (b) In TEM images, there is no visible effect of the treatment with GSH on the structures of the octahedra. Scale bar: 100 nm. (c) However, after the exposure to GSH, the bare octahedra are seemingly digested even faster in the presence of DNase I. While the untreated bare octahedra are fully degraded only after 24 h, there is no DNA origami band visible already after 2 h.

